# Imaging of MAP kinase dynamics reveals endocytic regulation of pulsatile signalling and network re-wiring in response to targeted therapy in EGFR-mutant non-small cell lung cancer

**DOI:** 10.1101/2024.05.14.594112

**Authors:** Alix Le Marois, Sasha Bailey, Steven Hooper, Sunil Kumar, Hugh Sparks, Yuriy Alexandrov, Deborah Caswell, Fabian Frӧhlich, Karin Schlegelmilch, Karishma Valand, Matthew Martin, Ana Narvaez, Charles Swanton, Julian Downward, Christopher Dunsby, Paul French, Erik Sahai

## Abstract

A better understanding of the signalling mechanisms underlying transitions from drug-sensitive to drug-tolerant states is required to overcome therapy failure. We combined single-cell biosensor imaging with functional perturbations to investigate the regulation of oncogenic signalling in EGFR-mutant lung adenocarcinoma. We find that despite the constant presence of the mutant oncogene, ERK signalling exhibits pulsatile dynamics, with pulse characteristics determined by the endocytic machinery. Analysis of drug-tolerant persisters (DTPs) revealed that, after an initial phase of complete pathway shut-down, signalling was rewired leading to renewed ERK pulses that drive cell cycle progression. FAK- and SRC-regulated adhesion complexes replace mutant EGFR as the driver of reactivated ERK pulses in DTPs, yet they remain controlled by the membrane trafficking machinery. We show that DTPs rely on additional survival pathways including YAP signalling, and that the phosphatase PTPRS represents a key node in therapy resistant cells, coordinating regulation of ERK, the cytoskeleton, and YAP.

## Introduction

Cancer is frequently driven by mutations that activate mitogenic signalling pathways, with EGFR L858R, EGFR exon 19 deletion, and KRAS codon 12 mutations frequently observed in lung adenocarcinoma (LUAD)^1,2^. These alterations promote ERK/MAPK and PI3K signalling^3,4^, which has led to the development of oncogene-specific inhibitors, with EGFR inhibitors such as osimertinib in use for over ten years^5,6^ and KRAS G12C inhibitors recently approved^4,7^. Regrettably, the development of resistance to EGFR inhibition is common, and typically occurs after several months of treatment, with cells described to be in a slow-growing, ‘drug-tolerant persister’ (DTP) state in the intervening period^8^. Recent studies have highlighted that the DTP state is itself heterogeneous in nature, prone to increased mutational rates^9^, with proliferative and non-proliferative sub-populations co-existing^10^. However, our understanding of how oncogenic signalling becomes rewired as cells transition through drug-sensitive, DTP, and finally to fully drug-resistant states is limited.

Activation of EGFR, either by ligand binding or mutation, triggers a series of inter-connected signalling and regulatory events, ultimately leading to ERK and AKT activation^3^ (Fig. 1A). Active ERK also phosphorylates and modulates the function of numerous cytoplasmic substrates, including many involved in transducing the signal from EGFR to ERK. These events typically inhibit signal transduction and constitute a negative feedback loop operating in a timeframe of less than an hour^11^. In addition, the expression of negative regulators of EGFR to ERK signalling is increased, which constitutes a second mechanism of negative feedback with slower kinetics^11^. These mechanisms limit the duration of ERK activation following ligand binding to EGFR. A further level of EGFR regulation is provided by endocytic pathways. Ligand binding promotes the internalisation of EGFR on endosomes, typically via a clathrin-dependent mechanism. Endosomes can either return receptors to the plasma membrane, or target them for degradation, terminating the signal^12^. The fate of EGFR molecules following endocytosis is largely determined by ubiquitin ligases and de-ubiquitinating enzymes (DUBs)^13–15^. Our understanding of the interplay between mutant EGFR, its endocytic fate, and ERK activity is incomplete in drug naïve-cells and largely absent in DTPs.

**Figure 1:**
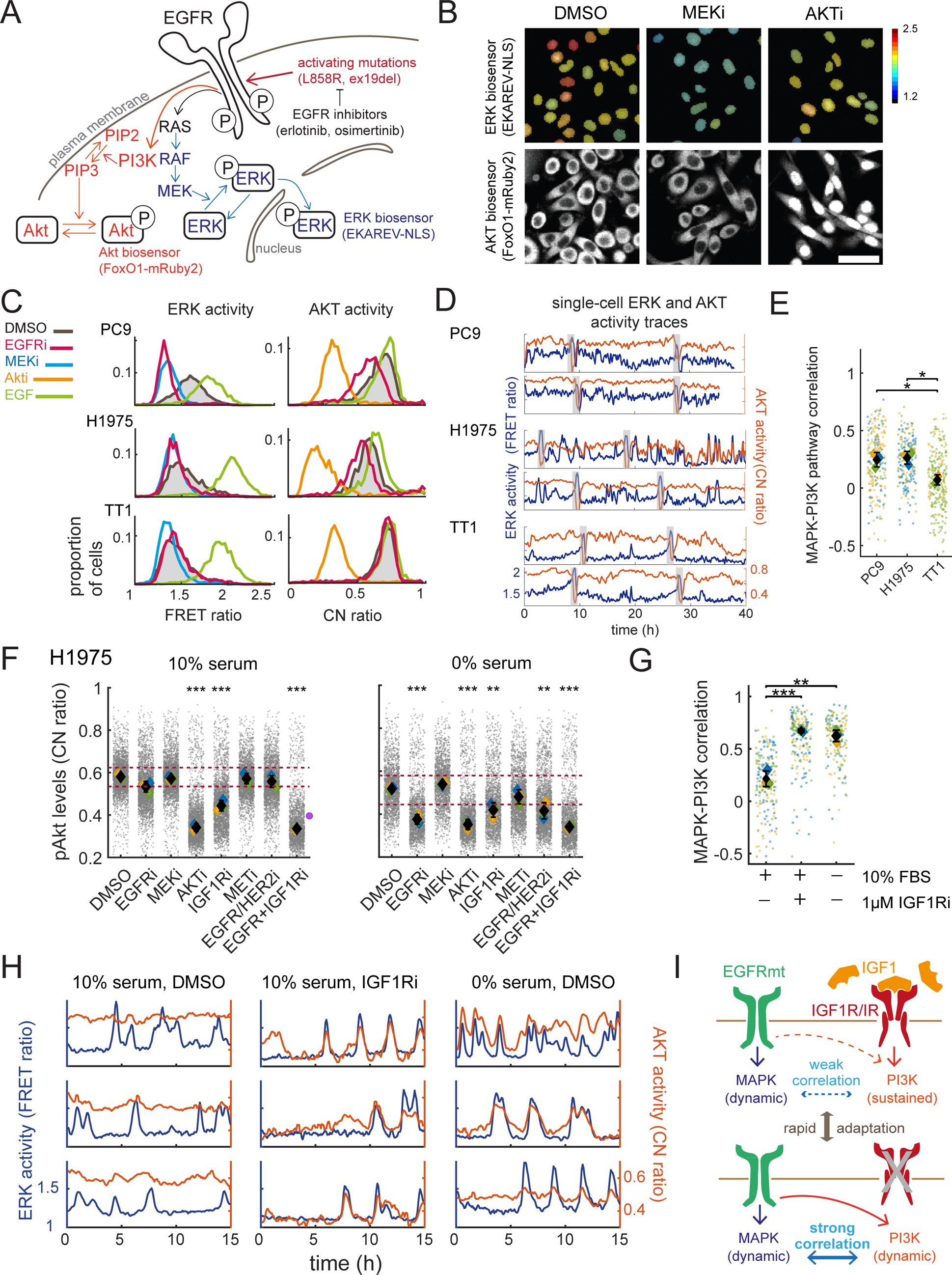
Imaging PI3K and MAPK pathway dynamics in EGFR-mutant NSCLC reveals dynamic patterns of oncogene-driven signalling. (A). Schematic of MAPK and PI3K pathway components downstream of mutant EGFR and the fluorescent biosensors used in this work. (B). Exemplar images of the EKAREV-FRET (top row, false colours) and the FoxO1-mRuby2 (bottom row) sensors in response to Akt or MEK inhibition. Scale bar is 50µm. (C). Distributions of ERK and Akt activity levels in PC9, H1975 and TT1 cell lines in response to MAPK and PI3K inhibition or stimulation. EGFRi: 500nM Osimertinib (PC9 and H1975) or 1µM erlotinib (TT1), MEKi: 100nM trametinib, AKTi: 1µM capivasertib, EGF: 20ng/mL. Data in (B) and (C) from N=3 independent experiments with n>1500 cells per condition. (D). Exemplar traces of pERK and pAkt levels in PC9, H1975 and TT1 cells tracked for a full cell cycle. Mitoses are highlighted by the grey rectangles. (E). Pearson’s correlation coefficients between the ERK and Akt traces in PC9, H1975 and TT1 cells. Each datapoint represents the correlation coefficient for an individual cell trace, black diamonds are the mean of the replicate medians. Data in (D) and (E) is from N=2 (TT1) or 3 (PC9 and H1975) independent experiments with a minimum of n=132 cells per cell line. (F). Akt levels in H1975 line in response to treatment with small molecule inhibitors in 10% (left) or 0% (right) FBS. Red dotted lines indicate the inter-quartile range of the DMSO control. Purple dot indicates the theoretical level of additive EGFR and IGF1R inhibition if the effects of both inhibitors on their own are added. DMSO: 0.1% v/v DMSO, EGFRi: 500nM Osimertinib, MEKi: 100nM trametinib, AKTi: 1µM capivasertib, IGF1Ri: 1µM NVP-AEW541, METi: 1µM savolitinib, EGFR/HER2i: 1µM lapatinib. Data is from N=2 (IGF1Ri+EGFRi) or N=3 independent experiments with a minimum of n=1000 cells per condition. (H). Pearson’s correlation coefficient between the ERK and Akt traces for individual H1975 cells in serum-free conditions or treated with IGF1Ri. Each datapoint represents the correlation coefficient for an individual cell trace, black diamonds are the mean of the replicate medians. (I). Exemplar ERK and Akt traces for untreated H1975 cells in 10% serum compared to untreated cells in 0% serum or cells treated with IGF1R inhibitor. Data in (H) and (I) is from N=3 independent experiments with a minimum of n=152 cells per condition. (J). Summary diagram of model for dynamics and coupling between MAPK and PI3K pathways in EGFR-mutant NSCLC. **Statistical tests:** (E) Unpaired Student’s t-test between the medians of the replicates. (F) & (G) 1-way ANOVA between the medians of the replicates with Tukey’s multiple comparison test. (H) Unpaired Student’s t-test between the medians of the replicates. *: p<0.05, **: p<0.01, ***: p<0.001.

One common characteristic of systems in which an activator drives negative feedback with a time delay is pulsatile or oscillatory behaviour of the signal output^11,16^. Indeed, pulsatile ERK signalling is a feature of many biological systems^17–19^, with previous studies exhibiting ERK pulses when wild-type EGFR is activated by low ligand levels^20,21^. However, it remains unclear whether pulsatile signalling occurs when EGFR is locked in an active state through oncogenic mutation, how it might be regulated, and how ERK dynamics change as cancer cells become tolerant to EGFR inhibition.

In this work, we explore the signalling dynamics downstream of oncogenic mutations, with the ultimate goal of uncovering vulnerabilities of the DTP state to provide better therapeutic options. Focusing on EGFR-mutant LUAD, we employ long-term time-lapse imaging of cells expressing fluorescent biosensors to quantify signalling outputs downstream of EGFR. This uncovers a close association between pulsatile ERK signalling and the emergence of drug-tolerant cells. By combining imaging of signalling dynamics with a genetic perturbation screen, we reveal how pulsatile signalling switches from being dependent on EGFR in drug-naïve cells to matrix adhesion signalling in DTPs, and we discover that the endocytic machinery plays a key role in both contexts. Lastly, our approach shows that DTPs rely on diverse survival mechanisms including YAP signalling, and we uncovered an unexpected role for receptor-type protein tyrosine phosphatase sigma (PTPRS) in supporting both MAPK and YAP-mediated signalling and survival in drug-tolerant cells.

## Results

### Highly dynamic ERK activity in EGFR mutant lung cancer

To investigate signalling dynamics and responses to targeted therapy in EGFR-mutant lung cancer cells, we employed fluorescent biosensors. Specifically, we used a FRET biosensor, EKAREV-NLS^22^, to monitor ERK activity, and a Kinase Translocation Reporter^23^, FoxO1-mRuby2 (hereafter termed AKT-KTR), to monitor AKT activity (Fig. S1A-B). Both biosensors were stably co-expressed in EGFR-mutant PC9 (EGFR ex19del) and H1975 (EGFR L858R) cells, as well as the immortalised non-tumorigenic lung epithelial line TT1^24^. As expected, MEK inhibition yielded lower FRET ratios, while AKT inhibition led to strong nuclear localisation of the AKT-KTR, indicative of low activity (Fig.1B). Time-lapse imaging confirmed the durability of the responses (Fig. S1C-D) and the validity of the readouts throughout the cell cycle, except during mitosis, which was previously reported to be CDK1-dependent^25^. We therefore excluded the 180 minutes prior to mitotic exit from all analyses (Fig. S1C-D and Methods).

First, we established distributions of ERK activity in control conditions, following EGF stimulation, and with pharmacological perturbation of EGFR, MEK, and AKT (Fig. 1C). As expected, the distributions of ERK activity were clearly separated following EGF stimulation and MEK inhibition. PC9 and H1975 cells had higher ERK activity than TT1 (Fig. 1C, left), and ERK activity in NSCLC cells was effectively abolished by inhibition of mutant EGFR using osimertinib, recapitulating the MEK-inhibited condition, while ERK activity in TT1 cells was not affected by wild-type EGFR inhibition using erlotinib. AKT activity exhibited a high baseline in all three cell types, which was clearly abrogated my AKT inhibition, but showed low sensitivity to EGFR stimulation or inhibition (Fig. 1C, right). To our surprise, we observed a clear overlap between control and MEK inhibitor-treated distributions in EGFR-mutant H1975 cells, suggesting that many cells had minimal levels of ERK activity, despite expressing an active mutant of EGFR (compare black and blue lines in Fig. 1C, left). Control and MEK inhibitor-treated distributions also overlapped for PC9 cells, though this was less pronounced. We confirmed the observation of low ERK activity in a large proportion of H1975 cells using immunofluorescence imaging (Fig. S1E).

To understand the distributions in pathway activity highlighted by population-level analysis, we performed long-term timelapse imaging and cell tracking of over 120 cells per cell line, to capture ERK and AKT activity throughout complete cell cycles (Fig. 1D, Supp. Video 1). TT1 had low ERK signalling with infrequent, defined pulses, while PC9 cells typically had persistently high ERK FRET signal with less pronounced pulses and H1975 cells exhibited dynamic ERK activity with frequent pulses. This suggests that the overlap in the distribution between control and MEK inhibitor-treated cells observed in Fig. 1C does not reflect distinct cell populations with stable and distinct ERK activity levels, but instead reflects highly dynamic ERK activity. In support of this, the ERK activity histograms from individual cell cycles closely resembled the full population histogram taken at a single time point (Fig. S1F). These analyses demonstrate that mutation of EGFR does not lead to a stable increase in ERK activity, but highly dynamic behaviour with variations between cell lines, and that the heterogeneity in ERK activity observed at any given point in time does not reflect inter-cellular heterogeneity, but is the result of the dynamic and pulsatile nature of signalling in all cells.

### Mutant EGFR can drive synchronous ERK and AKT pulses

Population level analysis indicated that EGFR was only weakly coupled to AKT activity in control conditions (compare black, red, and yellow lines in Fig. 1C, right). In addition, analysis of individual cell traces revealed low levels of correlation between ERK and AKT activity (Fig. 1E). Many receptor tyrosine kinases (RTKs) besides EGFR can activate AKT. A phospho-RTK array on H1975 cells revealed that EGFR, MET, ERBB2, ERBB3, IGF1-R and IR showed significant phosphorylation levels (Fig. S1G), and similar results have been published for PC9 cells^26^. Inhibition of IGF1R, but not MET or ERBB2, reduced AKT activity (Fig. 1F). Timelapse imaging performed in the presence of IGF1R inhibitor or in serum-free media, which lacks IGF1, revealed striking coordination in ERK and AKT pulses (Fig. 1G-H, Supp. Video 2). Combined inhibition of IGF1R and mutant EGFR, or EGFR inhibition in serum-free conditions completely abrogated AKT signalling, indicating that the pulses observed in the absence of IGF1R signalling depend on mutant EGFR (Figs. 1F, S1H). These data reveal that mutant EGFR is capable of driving synchronised pulses of ERK and AKT activity in a context-dependent manner (Fig. 1I). Moreover, the almost identical kinetics of pulse termination (Fig. 1J) suggest a common mechanism controlling ERK and AKT pulse duration, and argue against MAPK and PI3K-specific mechanisms of negative feedback.

### Characterisation of ERK pulses in EGFR mutant lung cancer

We sought to characterise pulsatile signalling in more detail, focusing our analysis on the dynamics of ERK activity, which shows greater dependence on mutant EGFR than AKT. We found that such dynamic patterns were a recurring feature of EGFR-mutant NSCLC, with pulsatile behaviour visible in three further cell lines expressing the EKAREV-FRET biosensor (Fig. S2A). We explored the nature of these dynamics, and identified at least two classes of ERK pulses (Fig. 2A): cell-intrinsic interphase pulses where spontaneous pulsing appeared unconnected to that of neighbouring cells; and collective interphase pulses, which propagated in concentric rings from a core cell or group of cells, as already described in the contexts of collective migration^17^, drug resistance^27^ and tissue homeostasis^19^. We developed a collective pulse detection algorithm inspired by published work^28^, and interestingly, we found that while the EGFR-mutant NSCLC cells had a higher frequency of cell-intrinsic pulses (Fig. 2B-C, Fig. S2B), the non-transformed, EGFR-wt TT1 line displayed a higher proportion of collective pulses than the cancer cell lines (Fig. 2D-E, Fig. S2C-D, Supp. Videos 3a-b). Morphological analysis indicated that in cells capable of collective pulses, cell-cell contacts were characterised by protrusions rich in actin fibres, which were absent in PC9 cells (Fig. S2E). Thus, oncogenic transformation is associated with an increase in cell-intrinsic pulsing and a concomitant loss of collective ERK pulsing.

**Figure 2:**
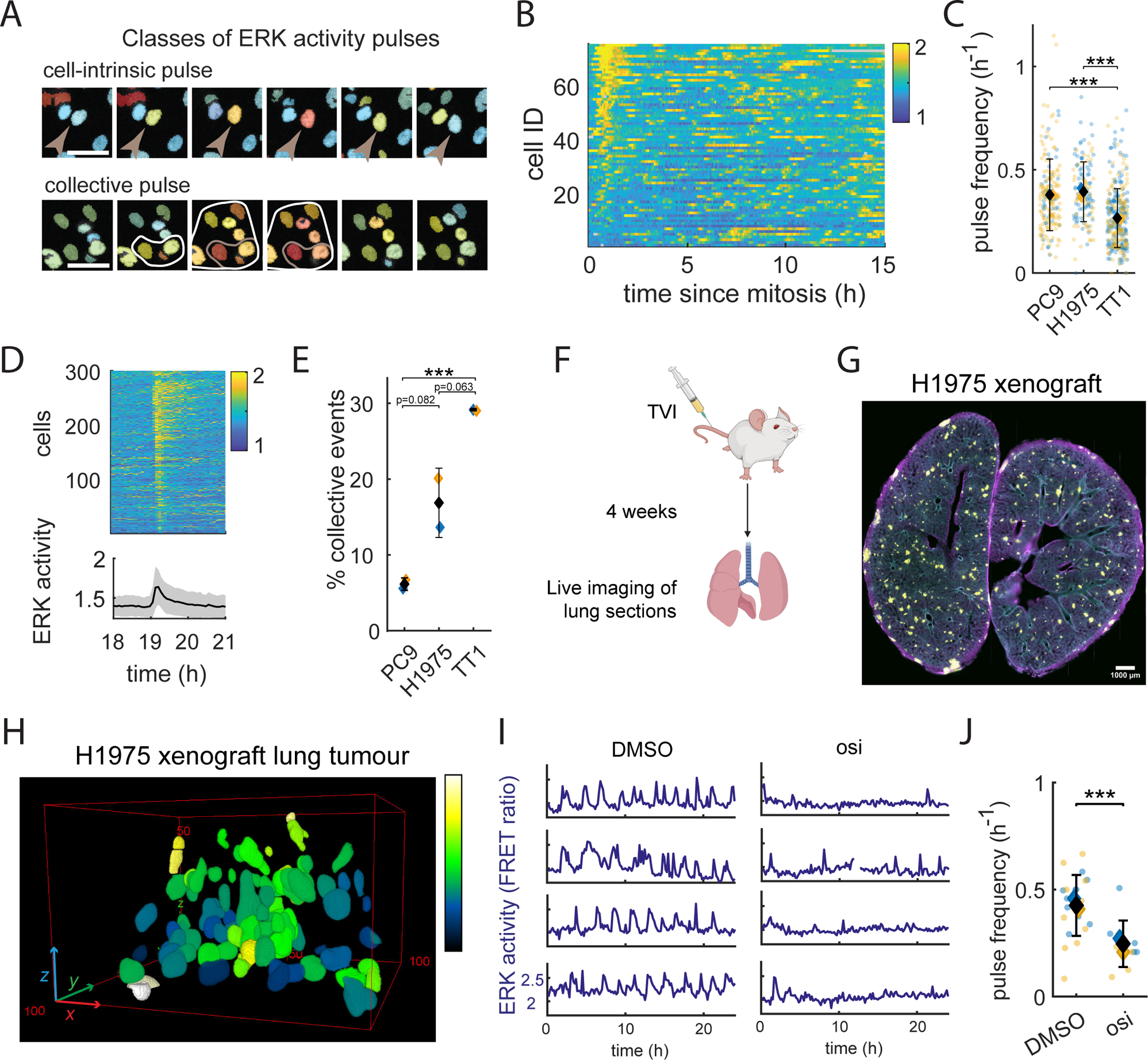
Characterising MAPK dynamics *in vitro* and *ex vivo* models of EGFR-mutant NSCLC. (A). Examples of cell-intrinsic and collective pulses of MAPK activity in the H1975 line. Scale bar is 50µm. (B). Kymograph of ERK activity in 70 H1975 cells aligned to mitotic exit. Pulses are visible as the yellow intervals throughout the interphase. (C). ERK pulse frequency in PC9, H1975 and TT1 cell lines. Data is from N=2 independent experiments and a minimum of n=132 cells per condition. Black diamond and error bar are mean + std of all data points. (D). Representative example of a collective pulse in the TT1 cell line involving approx. 300 cells. Above: Kymograph of the pulsing event, below: average trace with standard deviation in grey. (E). Proportion of positive ERK pulse events which are collective in PC9, H1975 and TT1 cell lines. Data is from N=2 independent experiments. (F). Schematic of experimental procedure to generate lung xenograft tumours. (G). Exemplar image of Precision-Cut Lung Slices of 2 lung lobes containing H1975 tumours. Representative of N=4 mice. Yellow: YFP from EKAREV sensor (tumours), Blue: DAPI (nuclei), Magenta: phalloidin (actin). (H). 3-dimensional rendering of a tumour within the lung epithelium acquired with OPM imaging of PCLS, with colour representing the EKAREV-FRET ratio. (I). Exemplar traces from H1975 cells in *ex vivo* tumours treated with DMSO or 500nM Osimertinib and imaged with OPM. Gaps in traces indicate mitotic events. (J). Frequency of ERK pulses in H1975 *ex vivo* tumours treated with DMSO or 500nm osimertinib and imaged with OPM. Data are pooled from 10 individual tumours from the lung tissue of N=2 mice per condition. **Statistical tests:** (C), (E) and (J): Unpaired Student’s t-test between the datapoints.

Next, we investigated ERK dynamics in the context of tumours growing within lung tissue. Orthotopic tumours were generated by tail-vein injection of H1975 or PC9-EKAREV cells into mice. Once tumours had formed, we generated Precision-Cut Lung Slices (PCLS) from tumour-bearing mice and performed imaging using a dual-view Oblique Plane Microscopy (dOPM)^29^ (Fig 2F-H, Fig. S3A). This revealed clear ERK pulses in H1975 tumours (Fig. 2I) that were strongly reduced with osimertinib treatment (Fig. 2I-J). Similar to observations in cell culture, pulses were present but less pronounced in PC9 tumours (Fig. S3, imaged with confocal microscopy). Together, these data establish pulsatile signalling as a recurrent feature in multiple models of EGFR mutant lung cancer, and demonstrate concordance between findings in cell culture and lung tissue contexts.

### Mapping MAPK and PI3K dynamics in the transition to drug resistance

Almost invariably, small populations of cells survive EGFR inhibition with osimertinib in patients, and this residual disease eventually leads to relapse. This can be modelled *in vitro*, where chronic exposure of drug-naïve (DN) PC9 cells to osimertinib leads to the emergence of a slow-growing Drug-Tolerant Persister (DTP) state^8^ after 1 week (Fig. S4A). H1975 cells undergo similar transitions, albeit with slightly slower kinetics. We characterised MAPK and PI3K activity in the transition from drug-naïve to drug-tolerant states. Upon first exposure to osimertinib, MAPK signalling was blocked in all cells with almost no pulses observed in either PC9 or H1975 cells (Figs. 3B, S4B). Longitudinal tracking of ERK activity and cell fate showed no difference in ERK activity between cells that died versus those that survived in the 24-72 hours following drug treatment (Fig. S4C&D), indicating that initial survival is not determined by variation in ERK activity. DTPs had elevated ERK signalling compared to DN cells under acute treatment, with pulsatile ERK signalling once again evident, but no longer dependent on mutant EGFR (Figs. 3A-D & S4B, Supp. Video 4). Abrogating the restored ERK pulses through MEK inhibition reduced cell viability (Fig. S4E). In contrast to PC9-DN cells, PC9-DTPs additionally exhibited collective pulses (Fig. S4F, Supp. Video 5). This was correlated with a change in cell morphology, with cell-cell contacts showing prominent F-actin fibres in PC9-DTPs (Fig. S4G). We additionally considered AKT activity, and found that it varied between cell lines, with low AKT activity in PC9-DTPs, and high activity in H1975-DTPs (Fig. S4H). Moreover, AKT activity was not pulsatile, and the two readouts were not correlated (Fig. S4I). Since the common feature in the two cell lines studied was a pulsatile reactivation of the MAPK pathway, our subsequent analysis concentrated solely on ERK regulation.

**Figure 3:**
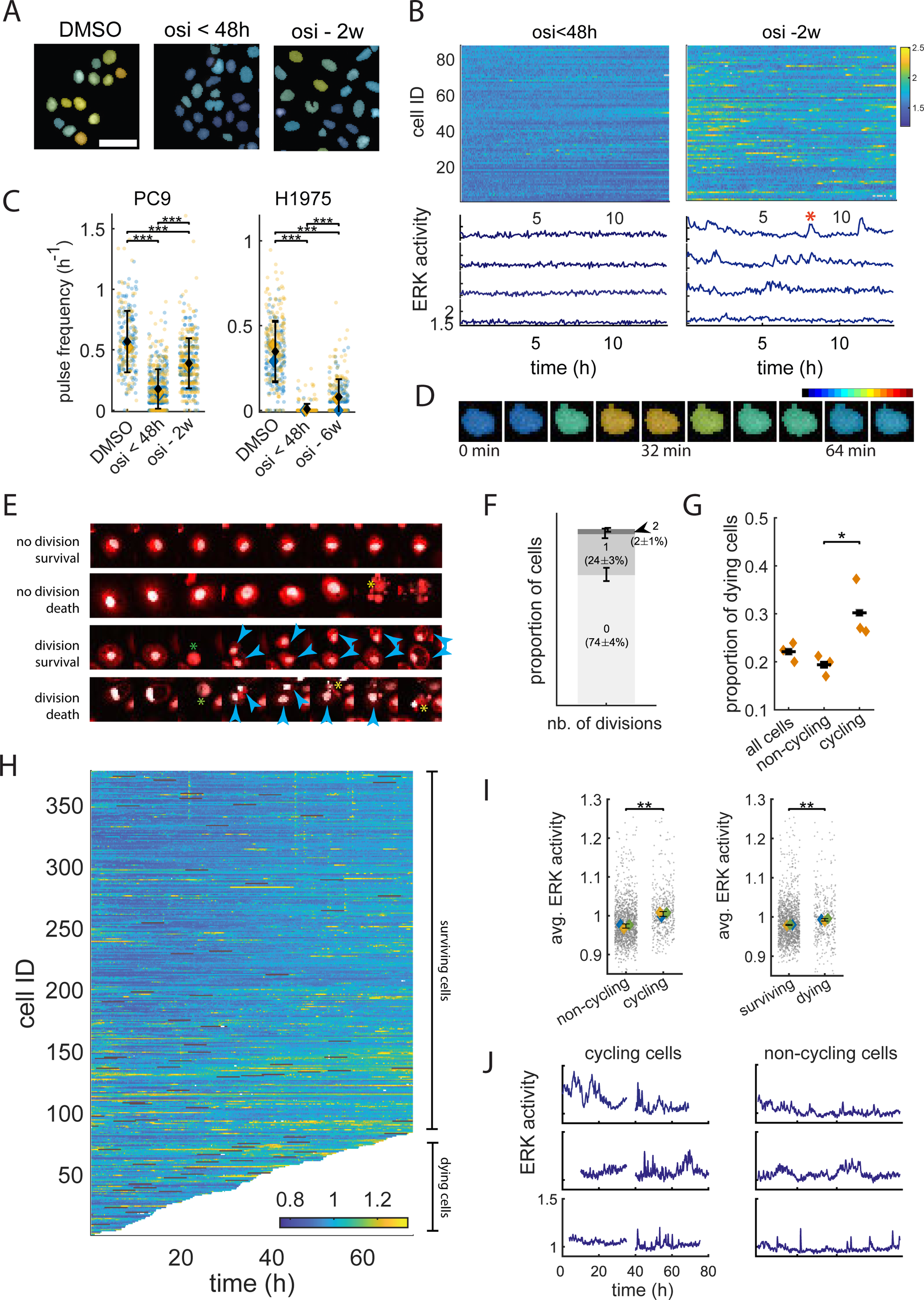
Partial re-activation of pulsatile MAPK activity drives cell fate diversification in drug-tolerant persister cells. (A). False colour FRET images of PC9 cells before and after treatment with 500nM Osimertinib. Scale bar is 50µm. (B). Kymographs of pERK traces and exemplar tracks for PC9 cells treated with Osimertinib for less than 48h (left) or 2 weeks (right). Data is representative of N>4 independent experiments. (C). Pulse frequency in PC9 (left) and H1975 cells upon acute or chronic osimertinib treatment. Data is from N=2 independent experiments with a minimum of n = 250 cells per condition. (D). False colour FRET image highlighting the ERK pulse in a PC9 DTP, highlighted by the red asterisk in Fig. 5(B). (E). Illustration of possible cell fate outcomes in PC9 DTP’s. (F). Break-down of the number of divisions within PC9-DTP populations observed for 72h. Data is from N=3 independent experiments. (G). Quantification of cell death events within non-cycling and cycling DTP populations. Data is from N=3 independent experiments. (H). Kymograph of pERK traces in PC9 DTP cells. Interrupted tracks highlight cells which dies within the 72h observation window. Mitoses are highlighted in red. Data is representative of N=3 independent experiments. (I). Quantification of ERK activity in PC9 DTP’s according to their cell fate. (J). Example ERK activity traces for cycling and non-cycling DTP’s. Gaps in traces indicate mitotic events. **Statistical tests:** (C). Unpaired Student’s t-test between individual data points. (G). Unpaired Student’s t-test. (I). Unpaired Student’s t-test between the medians of the replicates.

### Partial re-activation of MAPK activity is associated with cell cycle progression in DTPs

Unlike in the drug-naïve state where ERK was found to be active for some of the time in all cells, both PC9 and H1975-DTPs showed heterogeneity in MAPK activity, and some cells displayed very few or no ERK pulses (Figs. 3B & S4B). Heterogeneity in DTP proliferation and metabolic state has been highlighted recently^10^, and we hypothesised that MAPK activity might be a driver of cell fate divergence in DTPs. We tracked cell fates and ERK signalling levels in hundreds of individual PC9-DTPs for 72h (Fig 3E). The majority of tracked cells remained quiescent, with only 25% of traces containing 1 or more divisions (Fig. 3F), and cell death was observed in 23% of all traces (Fig. 3G-H). Intriguingly, the probability of cell death was significantly higher in the cycling DTPs than the non-cycling ones (Fig. 3G). Moreover, ERK activity was higher in both DTPs that cycled and those that died, compared to non-proliferating and surviving DTPs, respectively (Fig. 3I-J). These data suggest that the level of re-activated MAPK activity supports cell cycle progression but in doing so, also increases the risk of cell death. Given that MEK inhibitor treatment of established DTPs led to decreased cell numbers (Fig. S4E), and that after 4 weeks larger colonies of expanded DTP cells were observed, suggesting a net population growth (Fig. S4A), we conclude that MAPK reactivation supports cell fate diversification and overall population growth in DTPs (Supp. Table 1).

### The endocytic trafficking machinery is required for ERK pulses in drug-naïve cells

The analysis above indicates that, despite mutational activation, signalling is dynamic in mutant EGFR-driven lung cancer. Moreover, the emergence of DTPs is associated with restoration of pulsatile ERK signalling. Therefore, we sought to determine the mechanisms regulating pulsatile ERK signalling; first in drug-naïve cells and then in DTPs. To uncover the regulators of MAPK pulses, we curated an siRNA library targeting 90 genes expressed in H1975 cells^30–32^ known to interact with the EGFR-RAS-MAPK pathway, including EGFR ligands, receptor and non-receptor kinases and phosphatases, feedback inhibitors, and effectors of the membrane trafficking machinery (Fig. S5A-B). We performed 24h time-lapse imaging of H1975-EKAREV cells for each condition, using a custom-built high-throughput imaging platform (Fig. 4A). The data consisted of over 20,000 cell traces from 4 biological replicates, with a minimum duration of 6h. We generated 8 metrics for each trace related to temporal dynamics (pulse frequency, inter-peak distance), pulse shape (average pulse height, prominence, baseline, duration), as well as pulse-agnostic features (average and standard deviation of ERK activity levels)^21^. Principal Component Analysis (PCA) of these features showed that the variability of the data was well captured by 3 dimensions, with over 80% of total variance carried by the 3 first principal components (Fig. 4B). The variables related to temporal dynamics (pulse frequency and inter-peak distance) were almost orthogonal to a cluster of 3 highly correlated variables - pulse baseline, average pERK and pulse height (Fig. 4B). We therefore focused further analysis on the ‘pulse frequency’ and ‘pulse baseline’ variables (Fig. S5C), which captured most of the variation between different siRNA depletions. We used k-means clustering, maximising Dunn’s index with the condition that all controls should fall within the same cluster, to identify three clusters (Fig. 4C). Cluster 1 contained the control conditions and target genes with minimal effect on ERK dynamics (Fig. S6A), cluster 2 was characterised by low pulse frequency, and Cluster 3 by higher baselines but similar frequencies to Cluster 1 (Fig 4C-D, Fig. S6B-C).

**Figure 4:**
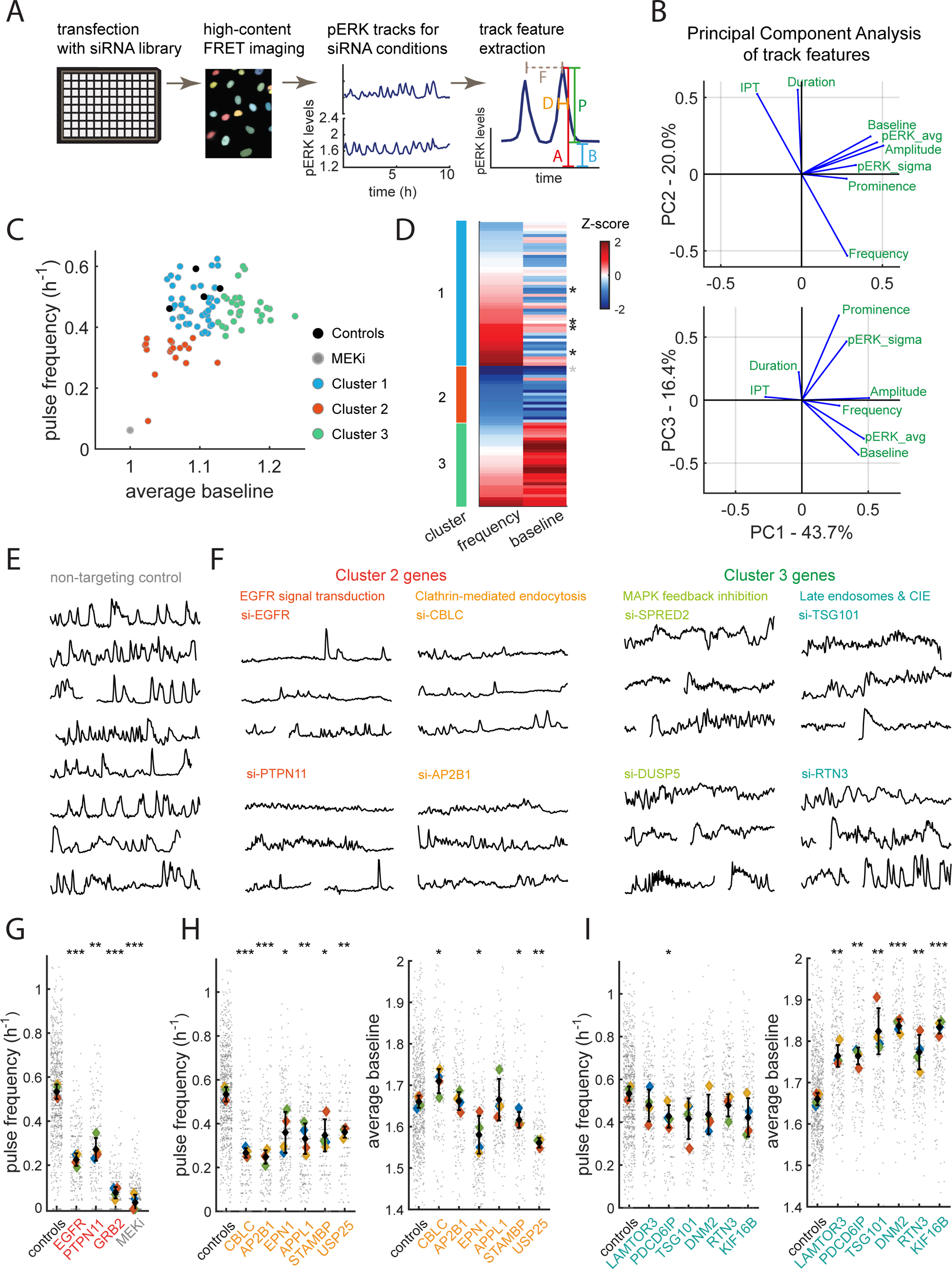
Live cell biosensor screen reveals central role of membrane trafficking machinery in regulating MAPK dynamics. (A). Diagram of live cell biosensor RNA interference screen workflow. (B). Representation of original trace variables in the space created by the PCA variables PC1-3. (C). Average baseline and pulse frequency for all screen targets. Colour of data points represents k-means clusters 1-3. (D). Frequency and baseline z-scores for all screen targets sorted by cluster. Negative and MEK-inhibited controls are shown by the black and grey asterisks, respectively. (E). Exemplar traces from the non-targeting control condition. Gaps in traces indicate mitotic events. (F). Exemplar traces from Clusters 2 and 3. Gaps in traces indicate mitotic events. (G). Pulse frequency from knock-down of Cluster 2 genes involved in EGFR-MAPK signal transduction. (H). Pulse frequency and baseline from knock-down of Cluster 2 membrane trafficking genes. (I). Pulse frequency and baseline from knock-down of Cluster 3 membrane trafficking genes. Data in this figure is from N=4 screen replicates, with a minimum of n=85 traces per condition **Statistical tests:** (G)-(H)-(I): unpaired Student’s t-test between replicate medians with FDR correction for multiple testing. *: p<0.05, **: p<0.01, *** : p<0.001.

Analysing the effect of individual gene depletions revealed both expected findings and new insights into the regulation of ERK pulses. In line with expectations, depletion of EGFR, GRB2 and PTPN11 (encoding the phosphatase SHP2) shifted cells into cluster 2 – low ERK pulses and low overall ERK activity (Fig. 4E-G, Supp. Table 2, Supp. Video 6). Depletion of negative regulators of EGFR-RAS-MAPK signalling (SPRED2, ERFFI1, PTPN1, DUSP5) increased ERK activity and pushed cells in cluster 3 – high baseline^33–35^ (Fig. 4E-F, Supp. Table 3, Supp. Video 6). More intriguingly, but not entirely unexpected, several molecules implicated in cytoskeletal regulation, including PTK2 and ITGA5 (encoding FAK and integrin α5), were also identified as positive regulators of pulsatile ERK activity (Supp. Table 2).

Our expectation was that perturbing the regulation and feedback inhibition of the EGFR-RAS-MAPK pathway would shorten or prolong individual ERK pulses. However, our data revealed that there was surprisingly little impact on this pulse characteristic throughout the screen. Indeed, the maximum effects of siRNA depletions on pulse duration were weaker than the effects on pulse frequency and baseline, and the pulse shapes did not differ strongly throughout the screen (Fig. S7A-B). This suggested that the control of pulse characteristics might be determined by other mechanisms. Given that endocytosis is both central to EGFR regulation and part of a cyclical process, we turned our attention to the effect of endocytic regulators on ERK dynamics. Molecules involved in clathrin-mediated endocytosis (CME) and early endosome formation, such as AP2B1, EPN1 and APPL1^36,37^, were positive regulators of ERK activity and required for pulse initiation (Fig. 4F&H, Fig. S6B). The depletion of several genes involved in guiding cargoes such as EGFR towards the late endosomal degradative pathway (TSG101, LAMTOR3 and PDCD6IP^12,37,38^), led to increased ERK baselines, such that pulsatile behaviour was less apparent (Fig. 4F&I, Fig. S6C, Supp. Video 6). Targeting of clathrin-independent endocytosis (CIE), which can also shuttle EGFR towards degradation, through knock-down of DNM2 (involved in both CME and CIE) and RTN3^39^, led to similarly elevated ERK activity (Fig. 4F&I, Fig. S6C). Lastly, depletion of KIF16B, which transports early endosomes towards the plasma membrane, also increased ERK baseline signalling, suggesting longer-lived early endosomes cause increased MAPK activity. Together, these data suggest that entry into the CME is necessary for pulse initiation, and that signal termination is linked to degradation. The reduction in pulses upon depletion of CBLC, STAMBP and USP25 support an important role of substrate ubiquitination in mediating EGFR-RAS-MAPK signal transduction (Fig. 4F&H, Fig. S6B, Supp. Video 6).

We further explored the effect of depleting effectors of membrane trafficking on EGFR levels (Fig. 5A, S8A). Blockade of CME through siRNA depletion of EPN1 or AP2B1 did not reduce EGFR levels, indicating that the reduction of ERK pulse frequency is not due to lack of EGFR (Fig. 5B, S8A). Blockade of late endosomes (TSG101, LAMTOR3), MVB degradation (PDCD6IP), CIE (DMN2 and RTN3), and anterograde transport of early endosomes (KIF16B) led to increased EGFR levels (Fig. 5C, S8A). Lastly, depletion of CLBC, STAMBP and USP25, which affect the ubiquitination of EGFR cargo and other proteins in the endocytic machinery, led to lower EGFR levels (Fig. 5B, S8A). These analyses suggest a nuanced relationship between EGFR levels and ERK activity. Comparison of EGFR levels and pulse frequency or ERK baseline across all 90 conditions revealed a weak correlation between receptor levels and a slightly stronger correlation with ERK baseline (Fig. 5D). Consistent with this, a weak correlation was observed between EGFR and pERK levels in unperturbed cell populations (Fig. 5E-G). In both cases, cells with low EGFR had low ERK activity, but cells with high EGFR levels did not always have high ERK activity. Thus, high EGFR levels are a necessary but not sufficient condition to generate ERK pulses, and we propose that sub-cellular localisation of EGFR within the endocytic machinery may control its ability to signal to the MAPK pathway. Co-localisation experiments showed that only a minority of intracellular vesicles loaded with active EGFR were also positive for RAS (Fig. 5H), suggesting that the discrete nature of MAPK signals could be linked to the physical separation of mutant EGFR and its downstream effectors. Our data argue that in drug-naïve cells, mutant EGFR is largely incapable of signalling to ERK, especially when at the plasma membrane, with pulses of signalling initiated by clathrin-mediated endocytosis and terminated by receptor degradation pathways (Fig. 5I). The temporal characteristics of individual pulse events is ‘hard-wired’ by global cellular control of endocytic dynamics and hence the duration of the pulses was not strongly altered by the manipulations tested.

**Figure 5.**
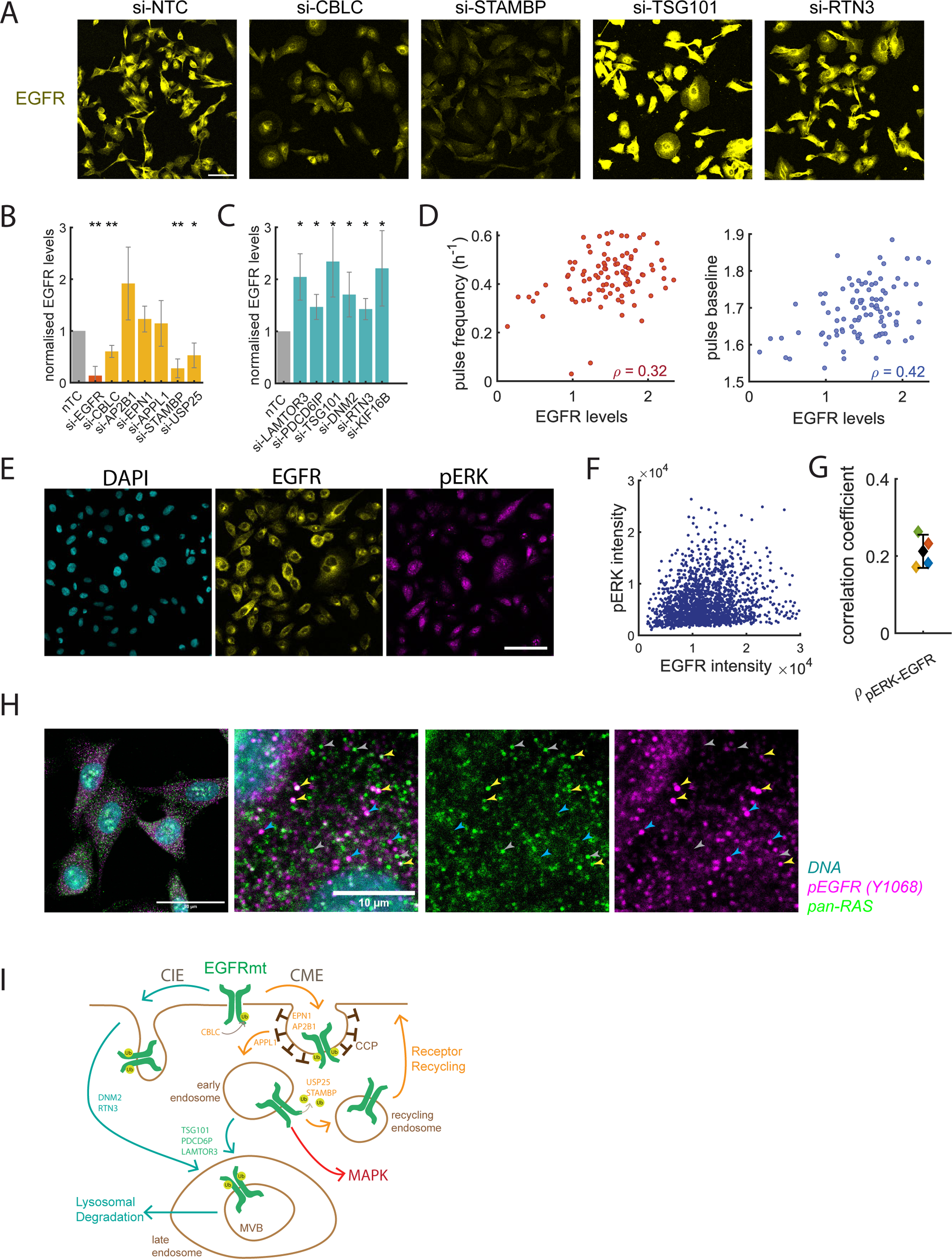
Elevated EGFR levels are a necessary but not sufficient condition to support MAPK dynamics. (A). Immunofluorescence imaging of EGFR in screen targets involved in membrane trafficking. Scale bar is 100µm (B). Quantification of EGFR levels in Cluster 2 membrane trafficking genes. (C). Quantification of EGFR levels in Cluster 3 membrane trafficking genes. Data in (A)-(C) is from N=4 independent experiments. (D). Correlation between EGFR levels and pulse frequency (left) and pulse baseline (right) from all screen targets. (E). Immunofluorescence imaging of EGFR and pERK in H1975 cells. Scale bar is 100µm. (F). Single cell correlation of EGFR and pERK signal intensities. Data is representative of N=4 independent experiments. (G). Correlation between EGFR and pERK signal intensity. Data is from N=4 independent experiments. (H). Confocal immunofluorescence imaging of RAS and phospho-EGFR in H1975 cells. Yellow arrows indicate vesicles positive for both markers, grey arrows are positive only for RAS and blue arrows positive only for phospho-EGFR. Data is representative of N=2 independent experiments. Scale bar is 50µM for full image and 10µm for zoomed-in insets. (I). Model showing the role of the membrane trafficking machinery in driving dynamic patterns of MAPK activity. CME: Clathrhin-Mediated Endocytosis. CIE: Clathrin-Independent Endocytosis. CCP: Clathrin-Coated Pits. **Statistical tests:** (B)-(C): unpaired Student’s t-test with FDR correction for multiple testing among the 95 screen targets.

### Pulsatile ERK signalling in DTPs remains dependent on endocytosis

We next investigated the regulation of MAPK signalling in DTPs, and acquired over 8000 MAPK activity traces in H1975-DTPs. To enable straightforward visualisation of network rewiring between the drug-naïve and drug-tolerant conditions, we projected the data on a single dimension using the pulse frequency and baseline metrics (Fig. 6A, Fig. S9A-B). As expected, EGFR was no longer a strong positive regulator of MAPK dynamics in H1975-DTPs. We previously observed that endocytic pathways were critical for ERK pulses in drug naïve cells. Interestingly, initiators of CME and cargo ubiquitination remained critical for the initiation of ERK pulses (Fig. 6A-C). However, the role of genes involved in late endosomal compartments was diminished compared to the drug-naïve case: only TSG101 depletion caused a trend towards higher pulsing frequency, and effectors of CIE no longer negatively regulated pulse dynamics (Fig. 6A, Fig. S9C-D). Therefore, we conclude that ERK pulses remain partially subject to regulation by the CME pathway in DTPs despite their independence from EGFR. In support of this hypothesis, we found that pulse duration was not significantly altered in DTPs cells compared to drug-naïve counterparts, suggesting that common mechanisms control pulse characteristics in both cell states (Fig. S9E), and a possible switching of key endocytic cargo from EGFR to other initiators of MAPK signals in the transition to DTP state.

**Figure 6:**
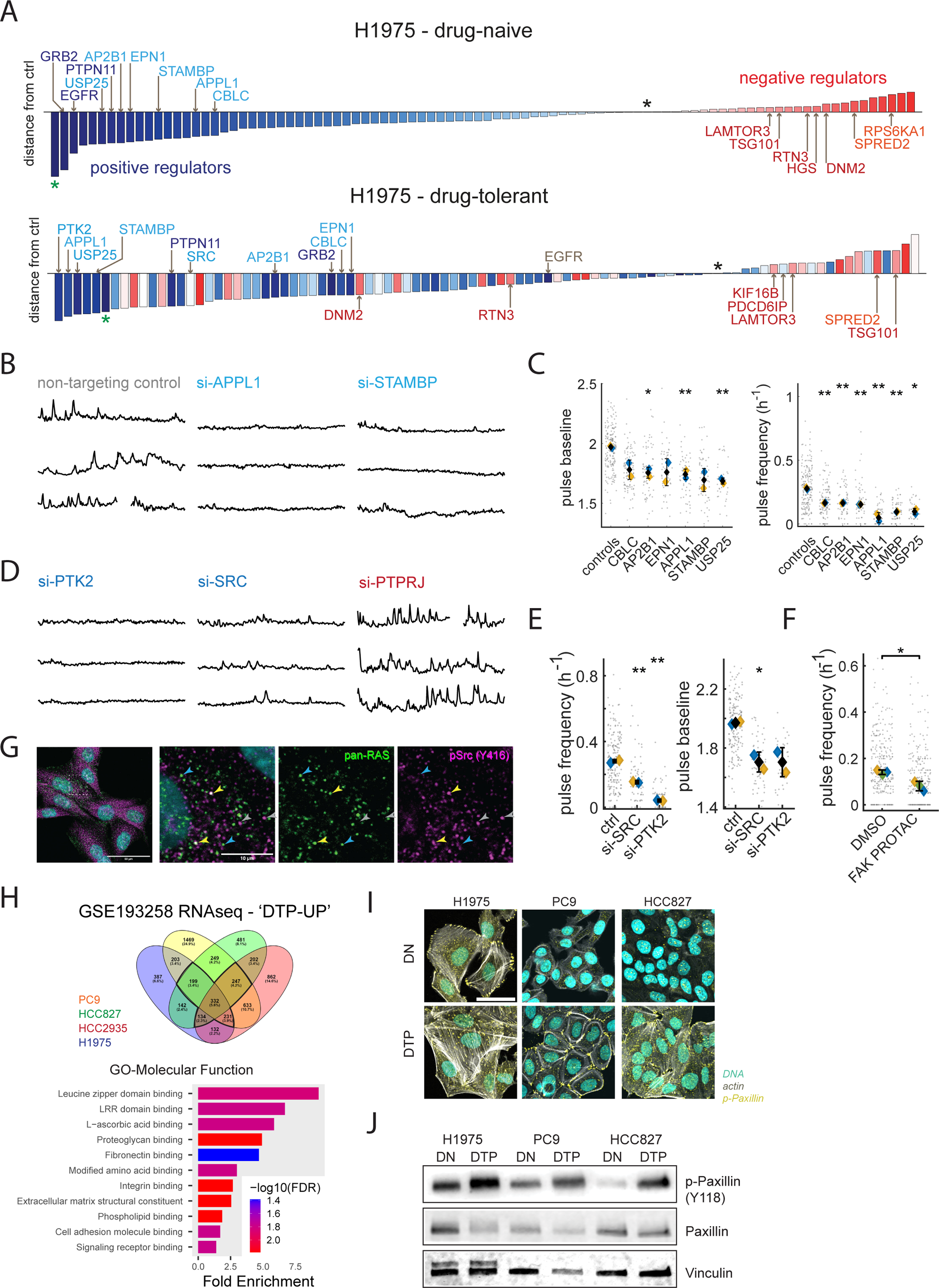
The DTP state is driven by re-wired cell adhesion-mediated signalling. (A). Bar charts of the screen targets between the drug-naïve and drug-tolerant states ranked in order of distance from the negative controls in terms of their effects on ERK pulse frequency and baseline. Bar colours from the drug-naïve conditions are conserved in the drug-tolerant conditions to highlight targets which are rewired. Black star indicates mean of controls, and green asterisk indicates MEK-inhibitor control. Data is from N=4 screen replicates in the drug naïve and N=2 replicates in the drug tolerant context. (B). Example ERK traces for some screen hits. Gaps in traces indicate mitotic events. (C). ERK pulse frequency and baseline with knock down of cluster 2 membrane trafficking genes. (D). Example ERK traces for some screen hits. Gaps in traces indicate mitotic events. (E). ERK pulse frequency and baseline with knock down of SRC (encoding for Src) and PTK2 (encoding for FAK) in H1975 DTP’s. (F). ERK pulse frequency in H1975-DTPs upon treatment with 1µM FAK PROTAC. Data is from N=3 independent experiments. (G). Confocal immunofluorescence imaging of RAS and phospho-Src in H1975-DTPs. Yellow arrows indicate vesicles positive for both markers, grey arrows are positive only for phospho-Src and blue arrows positive only for RAS. Data is representative of N=2 independent experiments. Scale bar is 50µM for full image and 10µm for zoomed-in insets. (H). ‘DTP-UP’ gene expression signatures based on bulk RNAseq data of drug-naïve and osi-DTP cell lines, with Gene Ontology analysis showing Molecular Function gene sets enriched in the DTP condition. (I) Immunofluorescence imaging of phospho-Paxillin pY118 and actin in osi-DTP cells. Data is from a minimum of N=2 independent experiments. (J). Immuno-blotting of phospho-Paxillin pY118 in drug-naïve and drug-tolerant cells. Data is from N=2 independent experiments. **Statistical tests:** (C), (E) & (F). Unpaired Student’s t-test between the replicate medians.

### A rewired adhesion-linked network drives ERK pulses in EGFRi resistant cells

Further analysis of our perturbation screen in DTPs revealed that PTK2 (FAK) depletion had the strongest effect on ERK activity in DTPs, with an increased role for Src additionally observed (Fig. 6A). Depletion of either PTK2 or SRC reduced ERK pulse frequency, while knock-down of negative Src regulator PTPRJ^40^ caused higher pulsing activity (Fig. 6D-E). SRC and FAK depletion decreased the viability of DTPs (Fig. S9F), albeit with the effect of SRC depletion limited to PC9 cells. We next tested the effect of pharmacological targeting of FAK and Src, using TT1 cells as non-transformed lungs cells that should remain viable. Consistent with previous work^41^, Src kinase inhibition using dasatinib repressed ERK signalling (Fig. S9G) and viability in DTPs (Fig. S9H), but was highly toxic to the TT1 cells^42–44^ (Fig. S9I). FAK inhibitors showed only moderate effects (Fig. S9J). We hypothesised that this might indicate a kinase-independent role for FAK. Therefore, we investigated reducing both kinase activity and protein levels of FAK using a PROteolysis TArgeting Chimera (PROTAC) approach^45,46^. Interestingly, treatment with a FAK PROTAC decreased the frequency of ERK pulses in H1975 DTPs (Fig. 6F), and reduced the viability of both drug-naïve and resistant cells (Fig S9J). Moreover, the FAK-PROTAC was well tolerated by TT1 cells at 1µM (Fig. S9I), suggesting that it may be a promising therapeutic strategy.

Given the prominent role of FAK and Src-family kinases in focal adhesion complexes^47^, these data are consistent with an increased role for cell-matrix adhesion in promoting pulsatile ERK signalling in DTPs. Interestingly, adhesion complexes share similar trafficking routes to EGFR, and can even be co-trafficked^48,49^, with additional evidence linking endocytosis to signalling from adhesion complexes to MAPK^50^. Accordingly, we found that a small number of intracellular vesicles were positive for both active Src and Ras (Fig. 6G). Gene Ontology analysis of a transcriptional signature for DTPs (‘DTP-UP’^51^), revealed that pathways related to cell-substrate interactions and production of extracellular matrix components were significantly enriched in DTPs (Fig. 6H), while integrin gene expression was significantly rewired (Fig. S9K). To investigate this in more detail, we probed drug-naïve cells and DTPs for actin and components of cell matrix adhesions. Fluorescence imaging revealed a denser network of dorsal and cortical actin fibres (Fig. 6I) and increased levels of focal adhesion protein phospho-paxillin, also seen by immuno-blotting (Fig. 6I-J), with its localisation at both focal adhesions and cell-cell contacts (Fig. 6I). Taken together, these results suggest that prolonged osimertinib treatment selects for cells with enhanced cell adhesion complexes that are coupled to ERK signalling via clathrin-dependent endocytic trafficking.

### PTPRS is a critical node in drug tolerant cells

Lastly, we interrogated our siRNA screen data to examine regulators which switched roles between drug-naïve and drug-tolerant states. Interestingly, many of the genes with a divergent role in ERK regulation between DN and DTP contexts were phosphatases (Fig. S10A). Of these, the membrane-spanning receptor-type tyrosine phosphatase sigma (PTPRS) caught our attention: it was required for ERK pulses in DTPs, but negatively regulated pulses in drug-naïve cells (Fig. 7A). PTPRS was included in the ‘DTP-UP’ signature (Fig. 7B), which we validated with immuno-blotting in three osimertinib-resistant cell lines (Fig. 7C). Depletion of PTPRS also reduced the viability of both PC9- and H1975-DTPs (Fig. 7D), and we confirmed these findings using a deconvolved siRNA strategy (Fig. S10B-E). Several studies have shown that PTPRS associates with endosomes^50,52^ and may interact with EGFR in this compartment. Accordingly, high resolution imaging revealed that PTPRS localised both at the plasma membrane and in intracellular vesicles, some of which were also positive for EEA1, a highly specific marker for early endosomes^53^ (Fig. 7E). We therefore propose that PTPRS modulates ERK pulsing activity in DTPs by associating with other endosomal cargoes.

**Figure 7:**
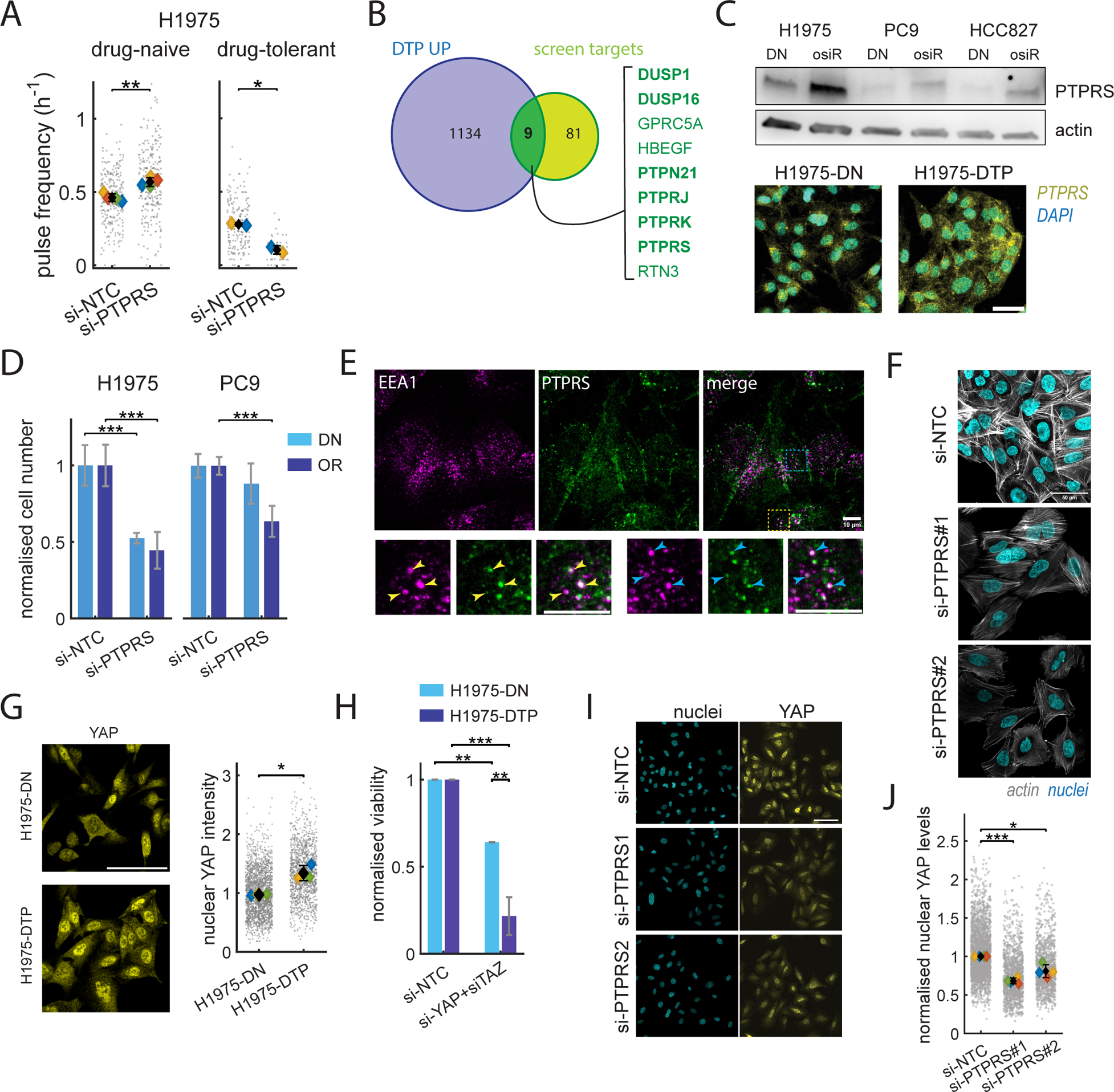
PTPRS as a bi-functional mediator of ERK signalling and survival in osi-DTPs. (A). Frequency of ERK pulses in drug-naïve and osi-DTP cells upon PTPRS knock-down. Data is from N=4 replicates in the drug naïve and N=2 replicates in the drug-tolerant contexts. (B). Overlap between the ‘DTP-UP’ gene signature and siRNA screen targets. (C). Immunoblotting of PTPRS levels in H1975, PC9 and HCC827 drug naïve and drug-resistant cells (above) and immunofluorescence imaging of PTPRS in H1975 cells (below). Data is representative of N=3 independent experiments. (D). Cell viability in H1975 and PC9 drug-naïve or osimertinib-resistant lines in response to PTPRS knock-down. Data is from N=3 independent experiments. (E). Confocal immunofluorescence imaging of EEA1 and PTPRS in H1975-DTP cells. Below: insets highlight areas of co-localisation. Scale bars are 10µm. (F). Confocal fluorescence imaging of actin in H1975-DTP cells upon PTPRS depletion. Data is from N=3 independent experiments. Scale bar is 50µm. (G). Immunofluorescence imaging of YAP1 in H1975 drug naïve and osi-DTPs with quantification of nuclear YAP intensity. Data is from N=3 independent experiments. Scale bar is 100µm. (H). Viability in H1975 drug naïve or osi-DTPs upon siRNA-mediated knock-down of YAP and TAZ. Data is from N=3 independent experiments. (I). Immunofluorescence imaging of YAP in H1975-DTP cells upon PTPRS knock-down. Data is representative of N=4 independent experiments. Scale bar is 100µm. (J). Quantification of nuclear YAP intensity in H1975-DTP cells upon PTPRS knock-down. Data is from N=4 independent experiments. **Statistical tests:** (A). Unpaired Student’s t-test on the replicate medians. N=4 for drug-naïve cells and N=2 for drug-tolerant cells. (D). Unpaired Student’s t-tests. Error bars show s.d. of the means. (G) & (J). Paired Student’s t-test on replicate means. (H). Paired Student’s t-tests. Error bars show s.d. of the means.

Further analysis indicated that depletion of PTPRS partially reverted the altered cytoskeletal organisation observed in DTPs, with a loss of F-actin fibres at cell-cell contact (Fig. 7F). Increased actin network density has been linked to regulation of the oncoprotein YAP, in turn activating a pro-survival transcriptional programme^54–56^. We therefore investigated YAP levels in DTPs, hypothesising that they might be modulated by the manipulations we identified previously. We confirmed that YAP activity, as judged by its nuclear accumulation, was increased in DTPs (Fig. 7G). Moreover, YAP and its homolog TAZ were necessary for DTP survival (Fig. 7H), and expression of an active YAP mutant^57^ was sufficient to confer some protection against osimertinib (Fig. S10F). FAK PROTAC treatment caused lower levels of nuclear YAP and decrease in actin signal intensity (Fig. S10G), further strengthening links between adhesion complexes, F-actin, and survival signalling. Interestingly, YAP levels in the nucleus of DTPs were reduced in the absence of PTPRS (Fig.7I). These results demonstrate that PTPRS is an unexpected bi-functional regulator of DTPs, being required for pulsatile ERK activity, and also for increased YAP1 functionality (Fig. 8).

**Figure 8:**
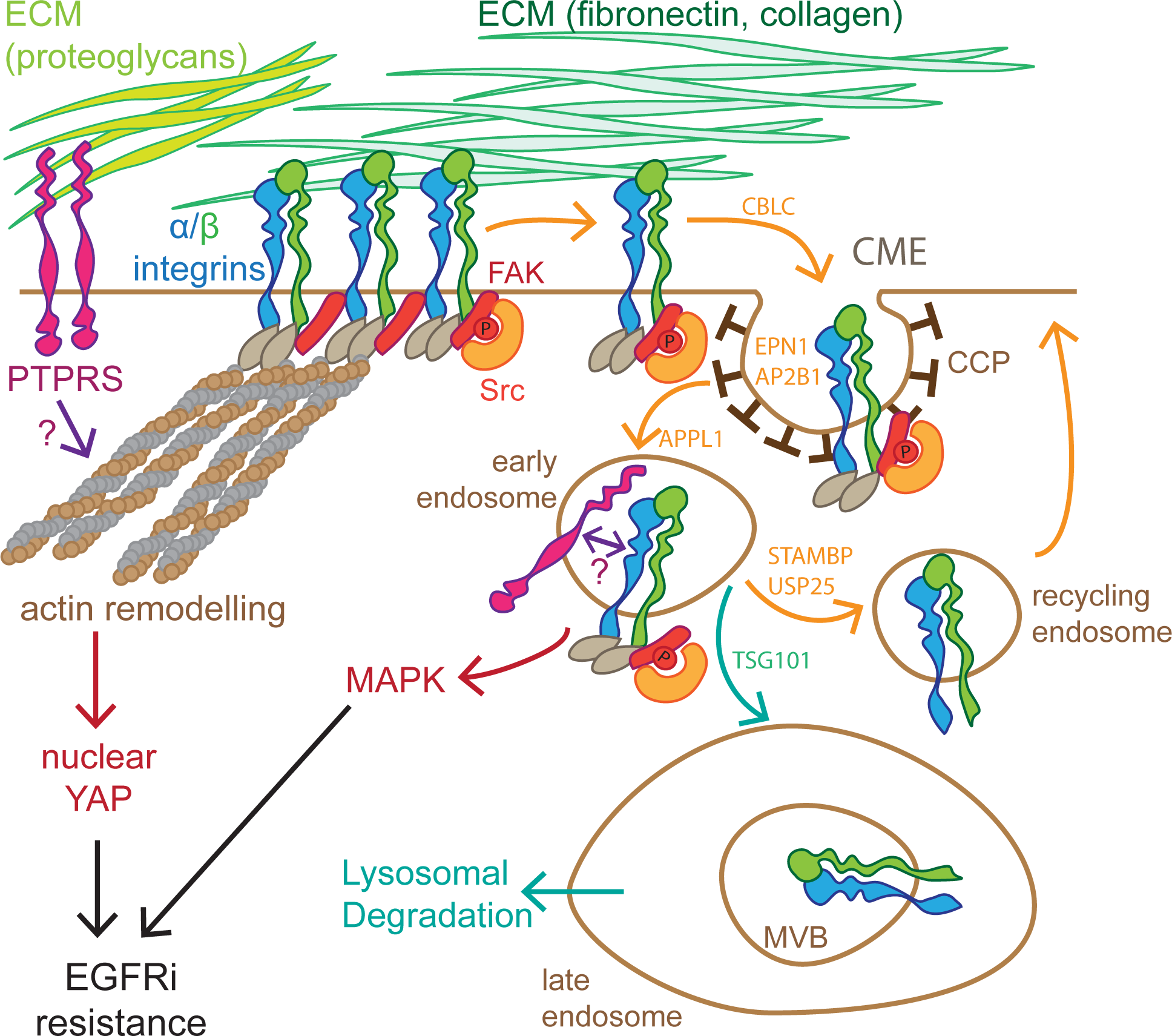
Model of rewired MAPK signalling and survival mechanisms in osi-DTPs. Abbreviations used: ECM (Extracellular Matrix), CCP (Clathrin-Coated Pits), CME (Clathrin-Mediated Endocytosis). ‘P’ indicates a phosphorylated residue.

## Discussion

Understanding why targeted kinase inhibition rarely leads to durable responses in patients requires knowledge of how the mutant kinase is coupled to its key downstream effectors, and how this coupling might evolve towards a loss of dependence on the oncogene. Given that cancer evolution acts at the level of the cell, an ability to study this process with single cell resolution is highly desirable. To address how signalling evolves during treatment with EGFR inhibitors, we established biosensor approaches to study long-term oncogenic signalling in individual cells.

Despite EGFR being mutationally activated, we found that ERK signalling is dynamic, with periods of inactivity interspersed with pulses lasting roughly 30 minutes. ERK pulses have been observed in various contexts, but the observation of pulsatile behaviour driven by a mutated receptor that does not require ligand binding for dimerisation and autophosphorylation is novel and surprising. It suggests that mutation alone is not sufficient to drive signalling, but that other events are required. Our analysis suggests that clathrin-mediated endocytosis of EGFR might be the trigger for signal initiation, and that the kinetics of cargo circulating through different components of the membrane trafficking machinery could determine pulse shape and duration. The relationship between endocytosis and wild-type EGFR signalling has been studied in some depth, with divergent findings on the precise nature and location of EGFR engaged in downstream signalling^58–62^. In the context of mutant EGFR, our data argue that although plasma membrane EGFR may be auto-phosphorylated, it is not engaged in signalling to ERK or AKT. We propose that the sporadic initiation the pulses is triggered by some stochastic event at the plasma membrane. Temporal fluctuations in higher order-EGFR complexes are a possible explanation: if a complex reaches a critical size or composition, then endocytosis and signalling are initiated. EGFR mutation favours this threshold being reached through a requirement for smaller complex sizes, but EGFR amplification could also yield similar behaviour by increasing the likelihood of large complexes. The requirement of mutant EGFR for collective ERK pulses, which others have reported to result from mechanical interactions between cells^17^, suggests that physical perturbation of the plasma membrane could be an additional mechanism for pulse initiation.

In many biological contexts, pulsatile dynamics result from the initiating signal promoting delayed negative feedback that ultimately terminates the pulse. Multiple mechanisms of negative feedback are induced by ERK signalling, including the expression of phosphatases. Our expectation had been that we would identify perturbations that affected pulse duration; however, this was not the case. Depletion of negative regulators, such as DUSP5 and SPRED2, elevated the baseline signal, but had minimal effect on pulse duration. Thus, instead of delayed negative feedback triggered by ERK signalling being the cause of pulse termination, we suggest that pulse termination and hence pulse duration, are determined by the dynamics of endocytic trafficking. Despite surveying a wide range of perturbations and cell lines, we only observed two dynamic regimes: one with a low baseline and ERK pulses of approx. 30 minutes at varying frequencies, and the other with a high baseline and less prominent pulses but of similar duration. The first regime was observed in H1975, EP2 and DTP lines (both PC9 and H1975), while the latter was found in PC9 cells and H1975 cells depleted for components of the endosomal degradation machinery. The HCC827 and H3255 lines displayed inter-cellular heterogeneity, with both regimes represented. Lastly, our observation of synchronised, EGFR-dependent ERK and AKT pulses when other inputs are removed, supports such a model, whereby signal initiation, duration and termination are regulated at the level of EGFR itself, rather than by independent negative feedback mechanisms acting further downstream, which would have different kinetics and refractory periods.

The transition to a drug-tolerant state was characterised by the almost total loss of ERK activity for a few days, before the re-emergence of pulsatile ERK signalling. Intriguingly, ERK signalling in DTPs was correlated with different cellular behaviours: cells with higher ERK activity exhibited both increased proliferation and increased cell death, whereas those with lower levels remained quiescent. The restored ERK pulses were no longer dependent on EGFR, but exhibited greater dependence on cell – ECM adhesion signalling. To our surprise, the duration of these pulses was similar to that of EGFR pulses in drug-naïve cells, and remained dependent on endocytic trafficking. We propose that EGFR inhibition selects for increased coupling of other cargoes trafficked through similar routes to ERK signalling. EGFR and other RTKs are known to co-traffic with integrins and other cell adhesion molecules which regulate downstream signalling, thus transitions in the relative coupling of these molecules could easily be selected.

Unfortunately, exploiting the membrane trafficking machinery as a therapeutic target will be difficult to reconcile with its central role in multiple normal cell functions. Hence, we focused on other more specific perturbations identified in our screen. The pro-survival function of PTPRS was initially surprising, as previous work has identified it as a tumour suppressor^63,64^ and a negative regulator of EGFR^65^. We propose that its switch to a positive regulator in DTPs is linked to the rewiring of upstream inputs into ERK away from RTKs and towards adhesion complexes, which is consistent with previous work in colorectal cancer^66^, and that PTPRS acts on endosomes to regulate the ability of cell adhesion-linked cargoes to activate MAPK signalling. Moreover, its additional role in YAP regulation renders it an important acquired vulnerability of drug-resistant cells. Specific substrates of PTPRS in this context remain to be identified, while its interaction with proteoglycans^67^, which are functionally enriched in DTPs (Fig. 6H), might explain the effects we observed on actin remodelling and YAP activity.

To conclude, the endocytic trafficking machinery is a critical regulator of pulsatile ERK signalling, even in cells with mutant EGFR. Network rewiring through the switching of endocytic cargo from EGFR to adhesion complexes and their associated kinases enables the partial re-activation of pulsatile MAPK activity and promotes the growth of drug-tolerant cells.

## Methods

### Cell Lines, plasmids and compounds

PC9 and H1975 parental cell lines were obtained from AstraZeneca. All other lines were obtained from the Cell Services platform at the Francis Crick Institute or generated by us^68^ (EP cell line, Fig. S2A). EKAREV-NLS cell lines were generated using the piggybac transposon system as described by us before^69^. Birefly, 2.10^5^ cells were plated in 6-well plates, and 48h later, were transfected with a 1:1 mix of pPBbsr2-EKAREV-NLS and pBase plasmids (4µg total DNA) and 10uL Lipofectamine 2000. Positive cells were expanded under blasticidin selection, and sorted using a BD Avalon flow sorter to select for cells with median expression levels of the biosensor.

The FoxO1-mRuby2 construct was cloned into the pLenti-puro lentiviral backbone using Gibson cloning and the pLenti-FoxO1-Clover^70^ and ERKKTR-mRuby constructs^23^ (Addgene plasmids nb 67759 & 90231, respectively). 1 million HEK293T cells were plated in 10cm dishes and transfected with the pLenti-FoxO1-mRuby2 and packaging plasmids VSV-G, GAG, and POL (16µg total DNA, 1:1:1:1 ratio). Media was replaced the following day, and viral production was promoted using 1mM sodium butyrate induction. On day 2, virus-containing medium was harvested and filtered through 0.45µm membranes, and added at 1:1 on target cells for infection. Infected cells were expanded under puromycin selection, and further purified using flow cytometry.

Cell lines were submitted to the Cell Services platform at the Francis Crick institute for authentication and Mycoplasma screening. Cells were sub-cultured in RPMI for a maximum of 12 weeks.

### Generation of drug-tolerant and drug-resistant cell lines

For DTP generation, PC9 cells were treated with 500nM Osimertinib for 2 weeks, H1975 cells were treated with 500nM Osimertinib for 6 weeks and passaged for a maximum of 6 weeks subsequently. HCC827 were initially treated with 50nM Osimertinib for 3 days, then 100nM for 6 days and finally 200nM for 6 days. Osimertinib-resistant (OR) lines were generated by treating cells with 500nM osimertinib continuously for 14 weeks. In all cases, osimertinib-containing media was replaced every 3-4 days.

### Live cell biosensor imaging: timelapse fluorescence microscopy

Cells were plated at approx. 2.10^4^/cm^2^ density in 24 or 96 wells plates (ibidi). Live imaging was performed 2 days after seeding.

The EKAREV FRET sensor was imaged using the ratiometric FRET method with laser-scanning confocal microscopy, as previously described by us^69^. Two systems were used throughout this project: an Olympus FV3000 microscope with a Plan-Apochromat 20x-0.75NA objective, 445nm laser excitation and 460-500nm and 530-630nm emission windows for CFP and YFP, respectively and a resolution of 1.24µm/pixel, or a Zeiss LSM710 microscope with a Plan-Apochromat 20x-0.8NA objective, 458nm laser excitation and 464-500nm and 518-555nm emission windows for CFP and YFP, respectively, and a digital zoom of 0.6 (1.38 µm pixel size). In both cases the pinhole was kept open for 2-D tissue culture imaging, and set to 1 Airy unit for optical sectioning in lung slice imaging experiments.

The FoxO1-mRuby2 sensor was imaged on the same fields of view as the FRET sensor, using a 561nm laser excitation and setting the pinhole to 3 Airy units.

Laser power was set to the minimum required to obtained a few saturated pixels in the image. Images were acquired every 4 to 6.5 minutes depending on the number of channels and fields of view per experiment. Photobleaching and phototoxicity were negligible in these conditions, with signal intensity remaining constant through the experiment, while population growth was exponential (∼18h doubling time) and virtually no cell death was observed in control conditions.

### Image analysis and Cell Tracking

#### Nuclear segmentation

A total intensity image was generated by adding the CFP and YFP images together. This was used to perform nuclear segmentation, either using ilastik (pixel annotation with 4 classes: background, nucleus, nucleus-background border and nucleus-nucleus border) or Stardist (fluorescent nuclei training dataset).

#### Biosensor quantification

Non-nuclear pixels in the image were classed as background, and average background intensity was subtracted from the raw images. Average CFP and YFP intensities were then computed for each nucleus, and the cell averaged FRET ratio was computed as: *FRET_ratio_ = YFP_av_/CFP_av_*.

For the Akt sensor, nuclear intensity *I_nuc_* was computed as the average intensity within a 5-by-5 pixel square centred around the centroid of the segmented nucleus. The nuclear mask was then subjected to a 2-pixel ring expansion, and the original mask was subtracted from the dilated one to generate a cytoplasmic ring mask. The brightest 50% pixels were used to compute the average cytoplasmic intensity *I_cyto_*. This was done to exclude background pixels from cells with an elongated morphology, where there were few cytoplasmic pixels on the sides of the nucleus. The CN ratio was then calculated as: *CN_ratio_* = *I_cyto_* / (*I_cnuc+_ I_cyto_*)

In addition to the biosensor values, 12 other metrics relating to nuclear morphology and signal intensity were calculated, using the ‘regionprops’ function in Matlab.

#### Cell tracking

The x,y,t coordinates of the segmented nuclei centroids were used to generate tracks with Trackmate^71^. A 25-pixel displacement threshold was used, and short tracks (<6h) were eliminated. Track data was then matched with biosensor and morphological data using custom-written Matlab scripts.

For most tracking cases, track splitting was not permitted as we found a large number of ‘false positive’ splits linked to segmentation errors or the presence of binucleated cells which did not correspond to mitotic events. However, for the siRNA screen (Figs. 4&5) and DTP cell fate tracking (Fig. 3), track splitting was permitted to maximise the number and length of tracks, and to preserve long-term cell identity. In these cases, track splits corresponding to true mitotic events were identified using a KNN learning algorithm trained on segmentation metrics from manually selected images of mitotic or non-mitotic events in the dataset. Splitting events which were within 2h of 3 consecutive frames identified as mitotic were flagged as true divisions, whilst all other splits were eliminated, conserving the longest track path. Remaining tracking errors mostly consisted in switches in cell identity across the track. To minimise these errors, tracks were manually reviewed, and tracks with clear discontinuities in nuclear area or FRET ratio were eliminated. Similar to previous reports^25,72^, we observed MEK-independent high FRET ratios and low (CN) ratios coincident with entry into mitosis. These events were easily identified by the abrupt change in nuclear area, caused by mitotic nuclear envelope breakdown. Using the correlated rapid increase in FRET ratio and nuclear area, mitotic exits were manually annotated, and the non MAPK-dependent FRET ratios in the 3h preceding mitotic exit were replaced with ‘NaN’ values.

### Immunofluorescence imaging and analysis

Cells were seeded as described for live cell imaging, and after drug or siRNA treatments, were fixed with 4% PFA for 20 minutes, rinsed twice in PBS, permeabilised with 0.1 Triton-X for 10 minutes and blocked for 1h with 10% serum (donkey or goat, depending on the species of the secondary antibody) in PBS+0.1% Tween-20 (PBST). Plates were then incubated with primary antibody dilutions in PBST overnight at 4°C, rinsed 3x in PBS and incubated with 1/500 secondary antibody dilutions and 0.5µg/mL DAPI in PBST for 1h at room temperature. For actin staining, phalloidin-Atto 633 (Sigma, 68825) was added at 20 nmol/L to the staining solution. Cells were then rinsed 3x with PBS before imaging using confocal microscopy.

### siRNA screen

#### Library preparation

The siRNA screen library was designed using the Horizon Discovery Cherry-Pick tool. 0.25nmol of siGenome siRNA smartpools were selected for the 90 targets and arranged in a 96-well plate format. Plates were centrifuged briefly, and each well was resuspended in 125uL of 5x siRNA suspension buffer reconstituted in RNAse-free water by thorough mixing. This generated 2µM siRNA suspensions. After gentle rocking at room temperature for 1h, the plate was aliquoted into 3 separate 96-well polypropylene plates and stored at −80°C.

#### siRNA transfection

siRNA was introduced into cells using a reverse transfection procedure. 40uL of Opti-Mem containing 1% Lipofectamine RNAiMAX was dispensed in each well of an ibidi 96-well plate. Then, 2.5µL of the 2µM siRNA screen plate was added and mixed into the transfection reagent mix. The plate was then left to incubate at room temperature for 20 minutes. Meanwhile, H1975-EKAREV-NLS cells were trypsinised and counted, and 10^4^ cells were added to each well in 200uL antibiotic-free, phenol red-free RPMI. This yielded a 20.6nmol/L siRNA concentration. There were 6 control wells, which included a non-targeting siRNA control pool (D-001206-14-05), an anti-GFP siRNA (P-002048-01-20), a MOCK transfection (siRNA reagent mix only), an untransfected well (Opti-MEM only), a MEK-inhibitor control (100nM trametinib) and an empty well with no cells for the background correction procedure. For the screen in drug-naïve cells, the medium was replaced with fresh RPMI 48h after transfection and cells were imaged 3h after media change. For the screen in drug-tolerant cells, cells were transfected in osimertinib-free media, which was replaced with fresh RPMI+500nM osimertinib the following day. As drug-tolerant cells grew more slowly, they were incubated for a further 48h before a second media change and imaging.

#### High throughput live-cell biosensor imaging platform

For imaging of siRNA-treated cells, a High-Content FLIM-FRET imaging platform previously designed by us^73^ based on a Olympus microscope body, was adapted for ratiometric fluorescence measurements in 96-well plate format and live cell imaging with home-built environmental control system maintaining a 37°C temperature and 5% CO2 atmosphere. CFP was excited using the blue band of a CoolLed pE-300 system (centred at 450nm) and a 20x/0.5NA objective (Olympus UPlan FL). CFP and YFP fluorescence signals were collected simultaneously using an Opto-Split II device (Cairn) fitted with a 505nm cut-off dichroic mirror and 480/40nm and 545/40nm emission filters, respectively, and imaged on a Retiga R1 CCD camera (QImaging). A single field of view was imaged for each well, and exposure time was set to 1s with a 6-minute acquisition frequency. This resulted in rectangular CFP and YFP channel images side-by-side within each frame. These raw images were registered and background-corrected using home-written algorithms using a reference-based, model-informed method for compensating XY- and time-dependent artifacts in time-lapse fluorescence microscopy data^74^ before subsequent segmentation, tracking and biosensor quantification as described above.

### *In vivo* experiments and Precision-Cut Lung Slice generation

The Francis Crick’s Institute Animal Welfare and Ethical Review Body and UK Home Office authority provided by Project License 0736231 approved all animal model procedures. Procedures described in this study were compliant with relevant ethical regulations regarding animal research. Lung tumours were generated by injecting 1.10^6^ cancer cells into the tail vein of 8-week-old NSG mice and left to grow for 4 weeks. The protocol published by Akram et al.^75^ was adapted to generate precision-cut slices of tumour-bearing lung tissue. 2% low-melting point agarose (Sigma, A9414) solutions were made in HBSS and kept at 42°C until use. The mice were sacrificed using overdose of anaesthetic (pentobarbital) via intraperitoneal injection. The thoracic cavity was then exposed; a cannula was introduced in the trachea and secured using a suture tie. 1mL of molten agarose was injected into the trachea to inflate the lungs and left to solidify on ice. The lungs were then removed and dissected into lobes. Lung lobes were then glued onto the platform of a Leica VTS-1200s vibratome, immersed in ice-cold HBSS, and sectioned into 300-µm thick slices. Slices were collected immediately and stored at 37°C and 5% CO2 in RPMI containing 10% FBS and 1% Penicillin-Streptomycin. Slices were imaged within 48h of sectioning.

### Dual-view oblique plane microscopy (dOPM) imaging and analysis

Dual-view oblique plane microscopy (dOPM) is a folded form of oblique plane microscopy (OPM)^76^. The original dOPM system reported in by us previously^29^ was modified here to use a 60×/1.2NA water immersion primary objective and a 50×/0.95NA air immersion secondary objective as described in^77^. Ratiometric FRET measurements were performed using a 445 nm laser for excitation, while CFP and YFP emissions were detected sequentially through 483/32 nm and 525/45 nm emission filters (Semrock, FF01-483/32 and FF01-525/45), respectively. Fields of view were imaged every 10 min for 24h. For each of the two dOPM views, z-stacks were acquired with planes spaced 1*μ*m apart and covering a scan range (in direction perpendicular to obliquely imaged plane) of 150*μ*m. Each image plane consisted of 512×512 pixels and the pixel size in sample space was 0.35*μ*m. The illumination light-sheet used had a calculated full width at half maximum of 3*μ*m at the waist in the sample plane. The two dOPM views were combined using the Multiview Fusion plugin available in ImageJ^78^. The details of this plugin’s use for dOPM are described in^29^. Registration information was obtained using 3*μ*m diameter fluorescence beads (Ultra-rainbow, URFP-30-2, Spherotech) suspended in aqueous 4% agarose gel. Following fusion, the datasets were converted to tiff stacks for segmentation with a 0.69 *μ*m pixel size in all three spatial dimensions. 3D segmentation of dOPM data was performed using a 3-scale version of nonlinear top-hat segmentation^79^ using 1.6, 3.2 and 6.4μm scale values^80^. The input (“nuclear intensity”) image *I_nuc_* for this detection was a linear combination of the CFP and YFP images *I_CFP_*, *I_YFP_*, weighted to equalize for signal-to noise ratio, namely as:

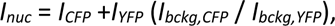

where *I_bckg,CFP_* and *I_bckg,YFP_* are background noise intensities estimated from corresponding global image histograms. The weights of the scale-smoothed images were 0.7/0.3 in favour of base scale, and the combined response threshold was set to 0.05. Post-thresholding steps included black and white morphology smoothing and hole filling, watershed clump separation, and small object and border objects removal. Segmentation quality was checked and tuned by visual inspection. FRET ratios were computed as described above.

### Survival Assays

Cells were plated in black-walled, clear-bottomed 96 well plates (Greiner, 655090). For siRNA treatments, the same transfection procedure as in the imaging screen was used with all volumes divided by 2 and 2500 cells seeded per well. Cells were then incubated for 72 or 96h post-siRNA transfection. For drug treatments, the medium was replaced 24h after plating with fresh RPMI containing the drug or equivalent DMSO control, and incubated for a further 72h. In all cases cell viability was quantified using the CellTiter Glo assay (Promega, G7571) according to supplier protocol, and luminescence was measured on a Tecan Infinite M1000 plate reader (1s integration time, no attenuation). For the osimertinib dose response experiments, survival data was fitted to a 3-parameter model as described by Nguyen et al.^81^, from which the efficacy was computed as the difference between maximum and minimum viability values obtained from the fitted parameters.

### Western Blotting

Cells were seeded in 10-cm dishes and treated with compounds as described in the previous sections. Plates were then placed on ice, rinsed with ice-cold PBS and lysed with 250uL RIPA buffer containing Phosphatase and Protease inhibitor cocktails (Roche PhosSTOP, 4906837001 and cOmplete, 11873580001). Lysates were then centrifuged at 13000rpm for 10 minutes at 4°C and supernatants were aliquoted and stored at −80°C. Protein content was determined using a BCA assay (Thermo Scientific, 23225), and sample absorbance was measured using a Tecan Spark plate reader. Laemmli buffer containing DTT was added to the samples, which were boiled at 95°C for 5 minutes before loading in pre-cast PAA gels (Bio-RAD, 4561084) and run at 100V for approx. 2 hours. Proteins were then transferred onto PVDF membranes (Bio-RAD, 1704156) using the Turbo-Blot transfer system. Membranes were then blocked using 5% milk in TBST, and incubated with 1/1000 antibody dilutions in TBST + 5% BSA overnight at 4°C. Membranes were then rinsed in TBST, and incubated with 1/10000 fluorophore or HRP-conjugated secondary antibody dilutions for 1h at room temperature. Luminescence and fluorescence signals were detected on Amersham ImageQuant 800 (Cytiva) and Licor-CLx digital imagers, respectively.

### Gene expression analysis

Gene expression data for the H1975 line was obtained from the MTAB2706^30^ and 2770 datasets^31,32^ as FPKM values on the EMBL-EBI website at https://www.ebi.ac.uk/gxa/experiments/E-MTAB-2706/Results and https://www.ebi.ac.uk/gxa/experiments/E-MTAB-2770/Results. The expression levels for genes of interest shown in Fig. S5B are the average of the two datasets, which were found to be highly correlated (ρ = 0.97 for the selected gene list).

The gene expression levels from the GSE193258 dataset were obtained from the Gene Expression Omnibus website in log2(TPM) format. For each condition (4 cell lines, drug-naïve and osi-DTP), the mean expression level was computed from the 3 replicates. The log2 fold-change (log2FC) was computed as the difference between the mean expression levels of DMSO and osi-DTP conditions. For each cell line, a threshold of 1 on log2TPM values was applied to select for genes expressed across the conditions, and a threshold of 0.6 on log2FC to select for genes with higher expression in the DTPs. The ‘DTP-UP’ signature was generated from genes found to be over-expressed in DTPs for 3 out of the 4 cell lines. Gene Ontology (GO) analysis of the ‘DTP-UP’ signature was performed using the ShinyGO online application^82^ (http://bioinformatics.sdstate.edu/go/), with the complete gene list from GSE193258 as background and the Molecular Function Gene Sets. Analysis of integrin expression levels was performed by extracting the list of all integrin genes from HUGO database (https://www.genenames.org/data/genegroup/#!/group/597 - 27 genes). Genes expressed above the threshold on average across all cell lines and conditions were then selected and fold change values were used to generate the heatmap.

### Data analysis and visualisation

Unless otherwise stated, all data visualisation and quantification was performed in Matlab (Natick, MA). Statistical testing was done using the ‘ttest’ (paired Student’s t-test), ‘ttest2’ (unpaired Student’s t-test) and ‘anova1’ (1-way ANOVA) functions. Correlation coefficients were calculated using the ‘corrcoeff’ function, and linear regressions were obtained using the ‘fitlm’ function. Images from microscopy experiments generated using ImageJ.

### Code and data availability

Data from the screen, Fiji scripts and Matlab code for biosensor processing and pulse analysis are available on GitHub at https://github.com/alixlemarois/dynamic_signalling. Other data from this study can be provided upon reasonable request to the corresponding authors.

**Figure S1.**
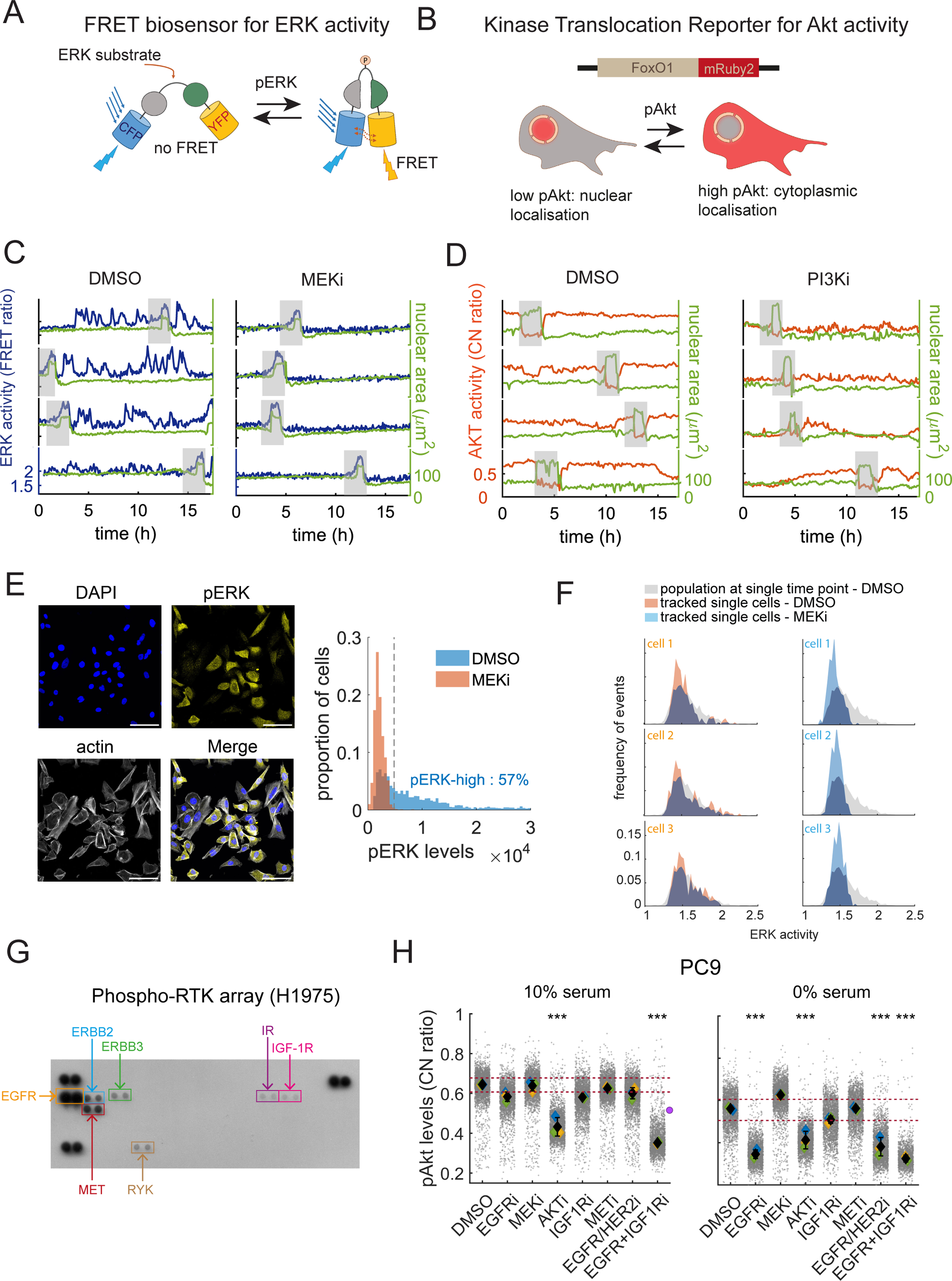
(related to Fig. 1) (A). Schematic of the EKAREV FRET biosensor to quantify ERK activity levels in individual cells. (B). Schematic of the FoxO1-mRuby AKT-KTR biosensor to quantify Akt activity levels in individual cells. (C). Exemplar traces of DMSO or MEK inhibitor-treated cells (100nM trametinib). ERK acitivity levels in blue and nuclear area in green. Red asterisks indicate mitotic events. (D). Exemplar traces of DMSO or PI3K inhibitor-treated cells (1µM pictilisib). Akt activity in orange and nuclear area in green. Grey rectangles indicate mitotic events. (E). Immunofluorescence staining of H1975 cells for pERK, counterstained with DAPI and phalloidin-Atto633. Scale bar is 100µm. Right: Distribution of pERK levels in DMSO or MEK inhibitor-treated cells as measured by fluorescence intensity levels of the pERK stain. Data is from N=3 independent experiments with a minimum of n=1900 cells per condition. (F). Analysis of the overlap between the distributions of ERK activity levels in single cells tracked for a full cell cycle, in control (left, orange histograms) or MEK-inhibited (right, blue histograms) conditions and in a large population of cells at a single time point (grey histogram). (G). Phospho-RTK array in H1975 cells. The main phosphorylated receptors are highlighted. Assay was performed once. (H). Akt activity in PC9 line in response to treatment with small molecule inhibitors in 10% (left) or 0% (right) FBS. Red dotted lines indicate the inter-quartile range of the DMSO control. Purple dot indicates the theoretical level of additive EGFR and IGF1R inhibition if the effects of both inhibitors on their own are added. See Fig. 1G legend for compounds used. Data is from N=2 (IGF1Ri+EGFRi) or N=3 independent experiments with a minimum of n=1000 cells per condition. **Statistical test**: (H) 1-way ANOVA between the medians of the replicates with Tukey’s multiple comparison test.

**Figure S2.**
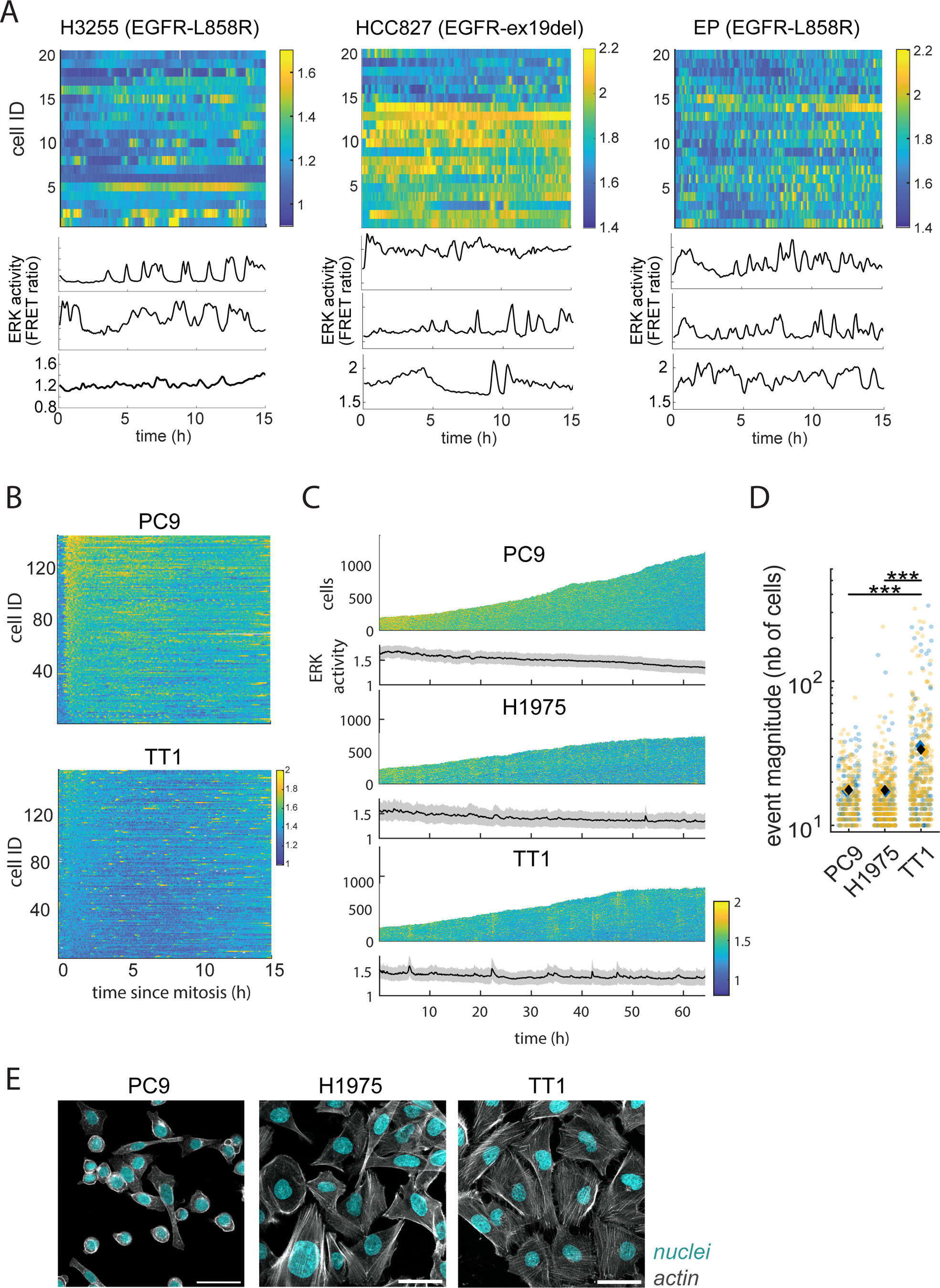
(related to Fig. 2) (A). Kymographs of ERK activity traces in 20 individual cells from 3 cell lines of EGFR-mutant NSCLC, with exemplar traces below. Data is representative of a minimum of N=2 independent experiments for each cell line. (B). Kymograph of ERK activity traces in PC9 and TT1 cells aligned to mitotic exit. Pulses are visible as the yellow intervals throughout the interphase. Data representative of N=3 independent experiments for each cell line. (C). Kymograph of ERK activity in time-lapse experiments for PC9, H1975 and TT1 cells, sorted by x-y position within the fields of view. The yellow vertical lines signify the presence of collective pulsing events, as also visible in the average traces below. (D). Magnitude of collective events for PC9, H1975 and TT1 lines. Statistical test: unpaired student’s t-test between the medians of the replicates. Data in (C) and (D) is from N=2 independent experiments. (E). Confocal fluorescence imaging of actin fibres in PC9, H1975 and TT1 cells. Data is from a minimum of N=2 independent experiments. Scale bar is 50µm.

**Figure S3.**
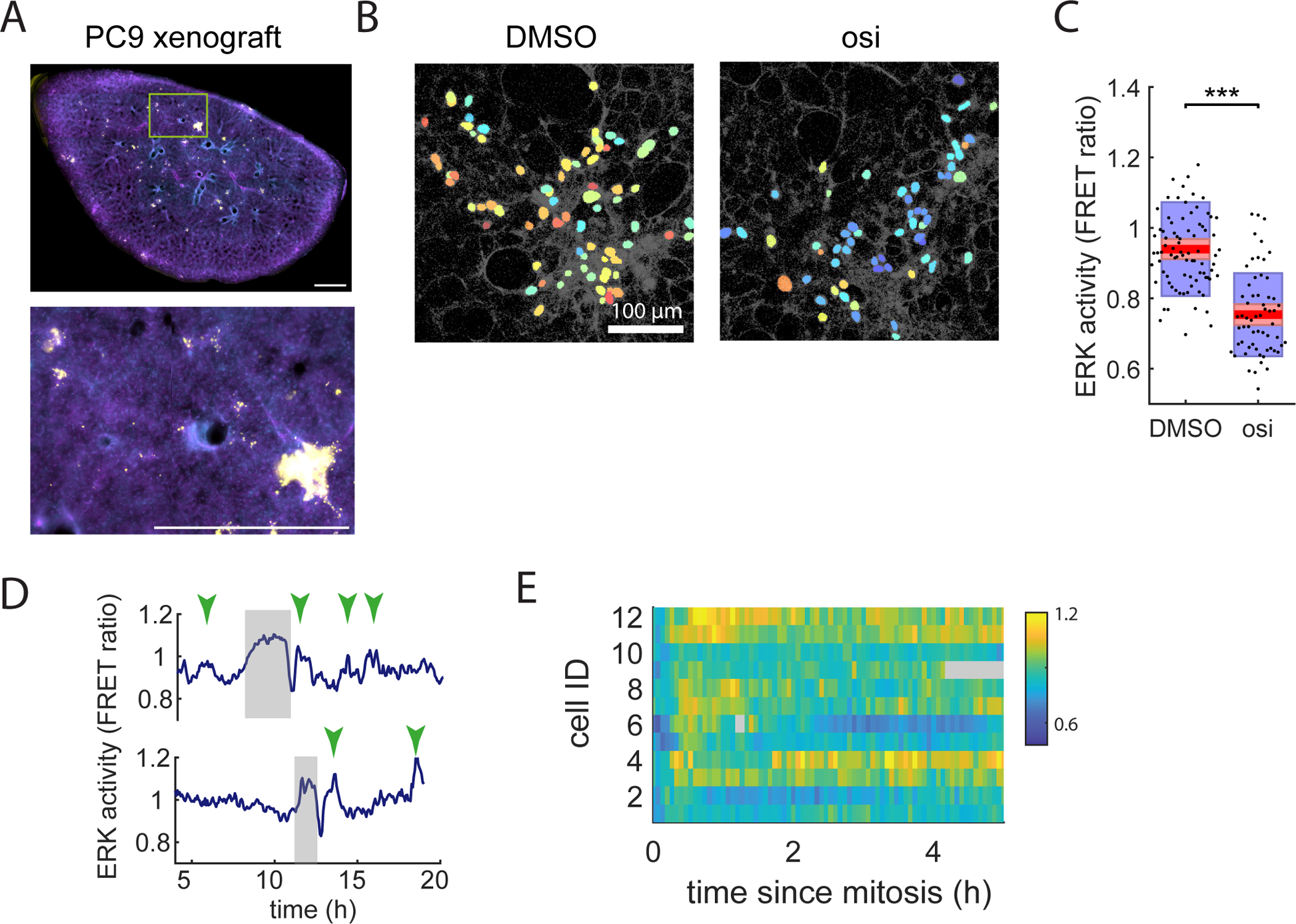
(Related to Fig. 2) (A). Exemplar image of Precision-Cut Lung Slice of a lung lobe containing PC9 tumours. Yellow: YFP from EKAREV sensor (tumours), Blue: DAPI (nuclei), Magenta: phalloidin (actin). Scale bar is 1mm. (B). Exemplar images of *ex vivo* PC9 tumours in PCLS treated with DMSO or 500nM Osimertinib. Tumour cells are in false colour reflecting their FRET ratio, while the surrounding lung epithelium is visible in grayscale. (C). Quantification of pERK levels in DMSO or Osimertinib-treated PC9 tumour cells in PCLS imaged by confocal microscopy. (D). Exemplar traces of PC9 cells in ex-vivo tumours, with mitotic events and pulses highlighted. (E). Kymograph of pERK traces in 12 individual PC9 cells in ex-vivo tumours, aligned to mitotic exit. Data in this figure is representative of lung slices and tumours acquired from a minimum of N=2 duplicate mice.

**Figure S4.**
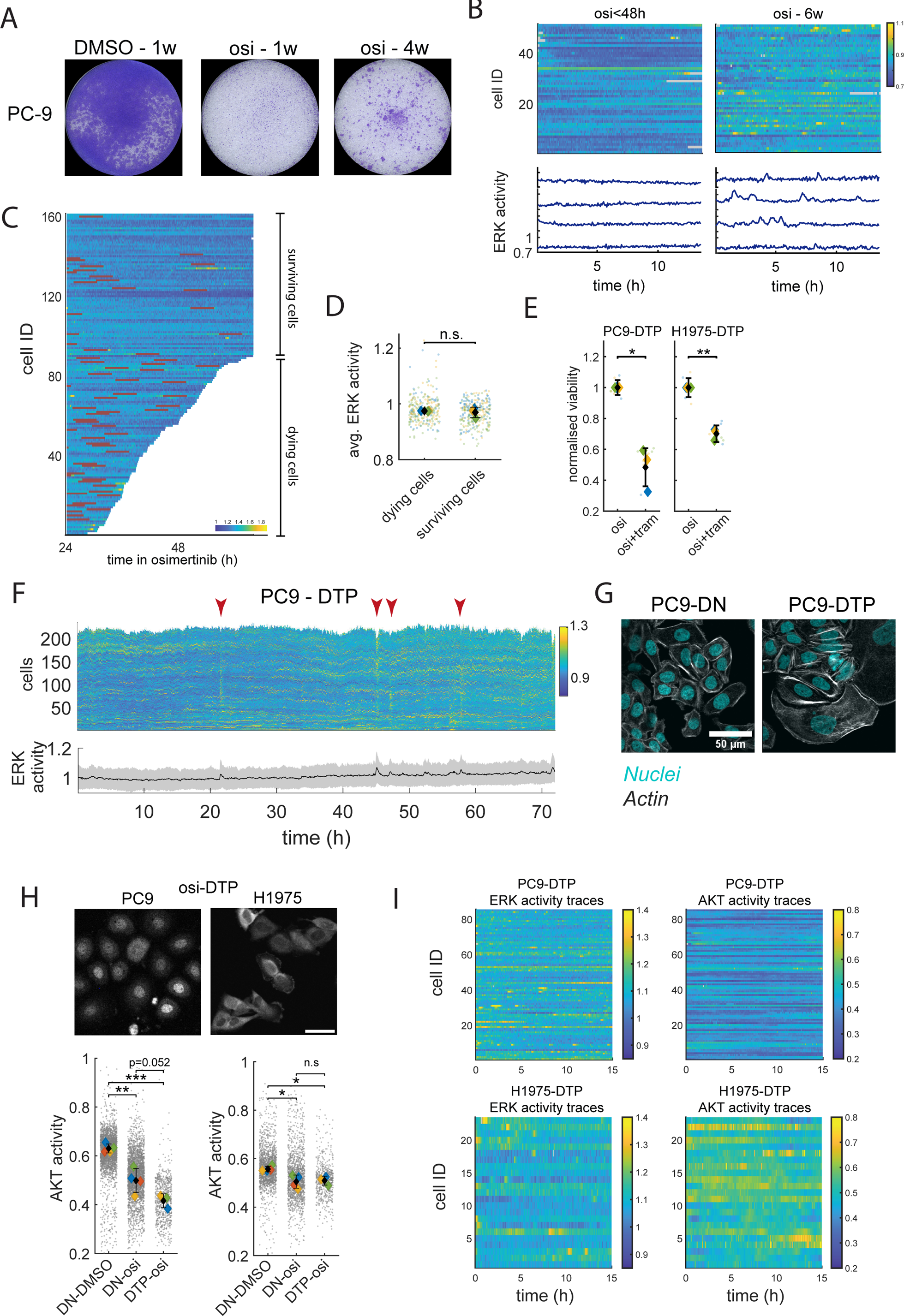
(Related to Fig. 3) (A). Crystal violet assay of PC9 cells treated with Osimertinib for 1 or 4 weeks. (B). Kymographs of ERK activity traces and exemplar tracks for H1975 cells treated with Osimertinib for less than 48h (left) or 6 weeks (right). Data is representative of N>3 independent experiments. (C). Kymograph of ERK activity traces in PC9 cells in response to acute Osimertinib treatment. Red bars indicate mitoses. Data is representative of N = 3 independent experiments. (D) Comparison of ERK activity levels between dying and surviving cell populations upon acute osimertinib treatment. (B). Effect of MEK inhibition on DTP viability. Data is from N=3 independent experiments each conducted in duplicate or triplicate. (C). Kymograph and average trace of ERK activity levels in PC9-DTPs illustrating the presence of collective pulses (highlighted by red arrows). (D). Fluorescence microscopy images of PC9 cells stained for actin. Images are representative of N=3 independent experiments. (E). Akt activity in PC9 and H1975 osi-DTP’s. Data is from a minimum of N=3 independent experiments per condition. Scale bar is 50µm. (F). Kymographs of ERK and Akt activity traces in the same cells in PC9 (above) and H1975 (below) osi-DTP’s. Data is representative of a minimum of N=3 independent experiments. **Statistical tests:** (D). Paired Student’s t-test between medians of the replicates. (E). Paired Student’s t-test between replicates. (H). Unpaired Student’s t-test between the medians of the replicates.

**Figure S5.**
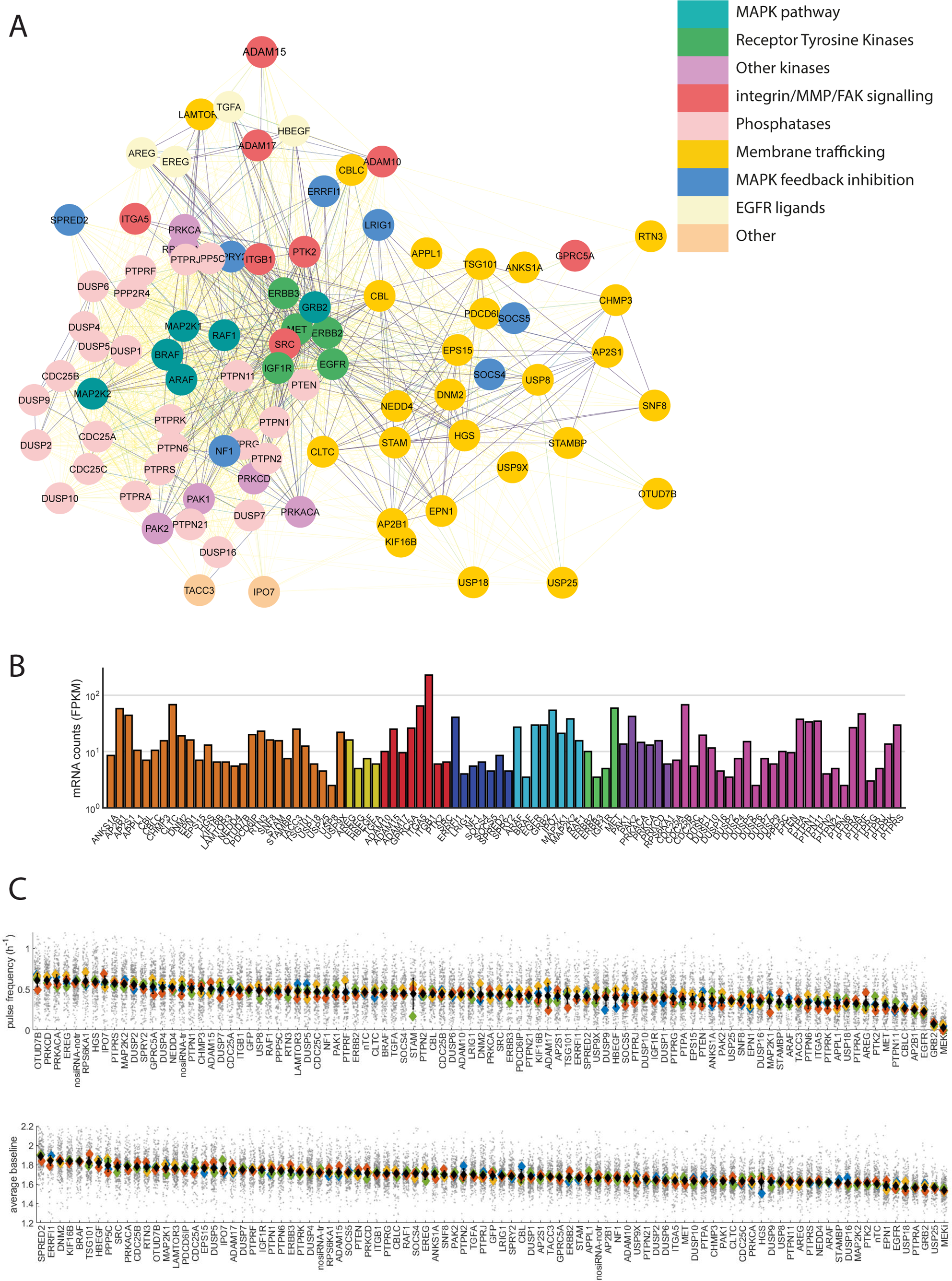
(Related to Fig. 4) (A). Network representation of the 90 screen targets. Darker lines between nodes signify stronger connections. Network connectivity data was obtained from the String database^83^ with the list of targets as input, and was visualised using the Cytoscape software^84^. (B). Expression levels of screen targets in H1975 cell line. (C). Pulse frequency and baseline upon siRNA mediated knock-down of all gene targets, sorted in decreasing order.

**Figure S6.**
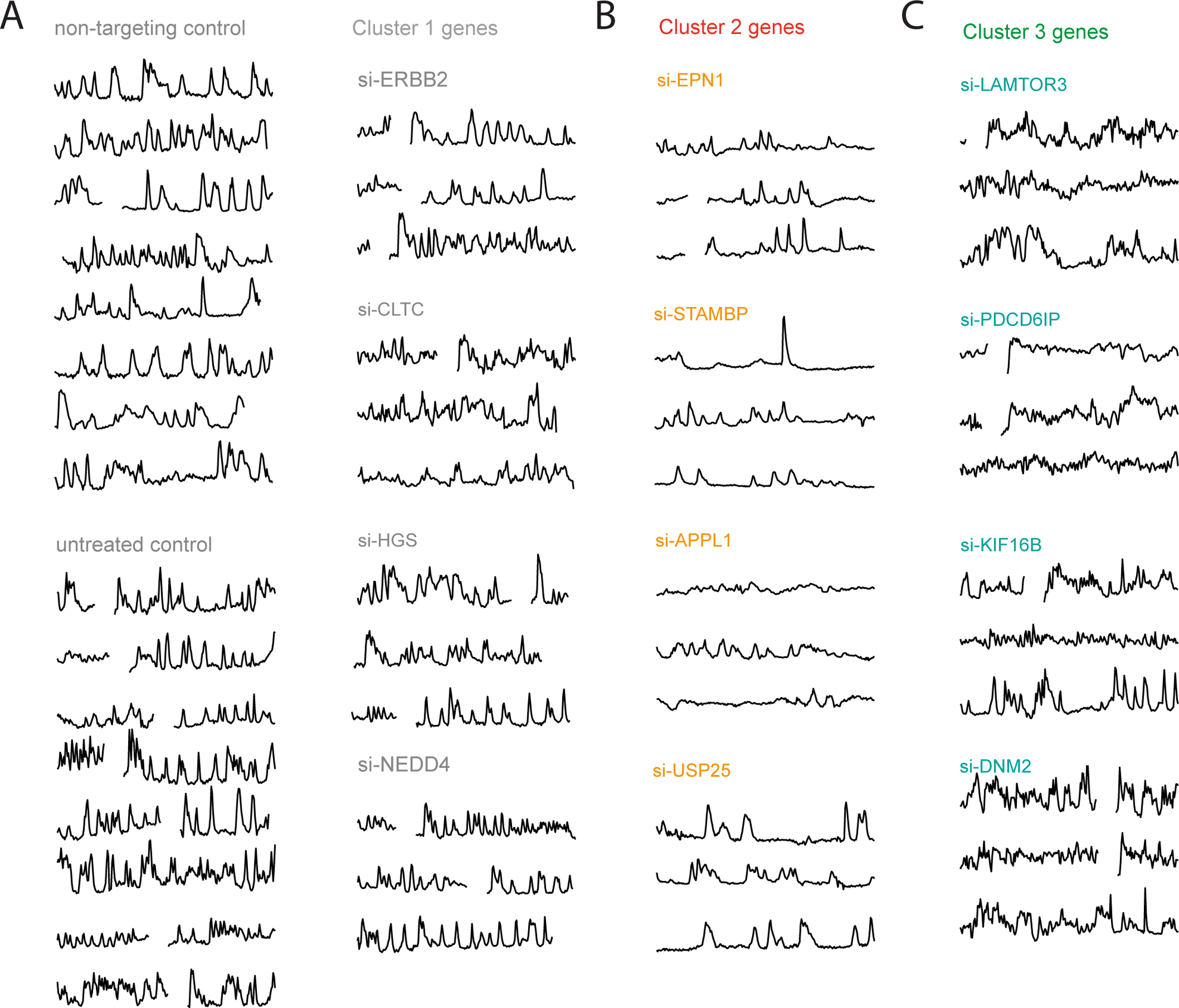
(Related to Fig. 4) (A). Exemplar tracks for controls and Cluster 1 genes. Gaps in traces indicate mitotic events. (B). Exemplar tracks for Cluster 2 membrane trafficking genes. Gaps in traces indicate mitotic events. (C). Exemplar tracks for Cluster 3 membrane trafficking genes. Gaps in traces indicate mitotic events.

**Figure S7.**
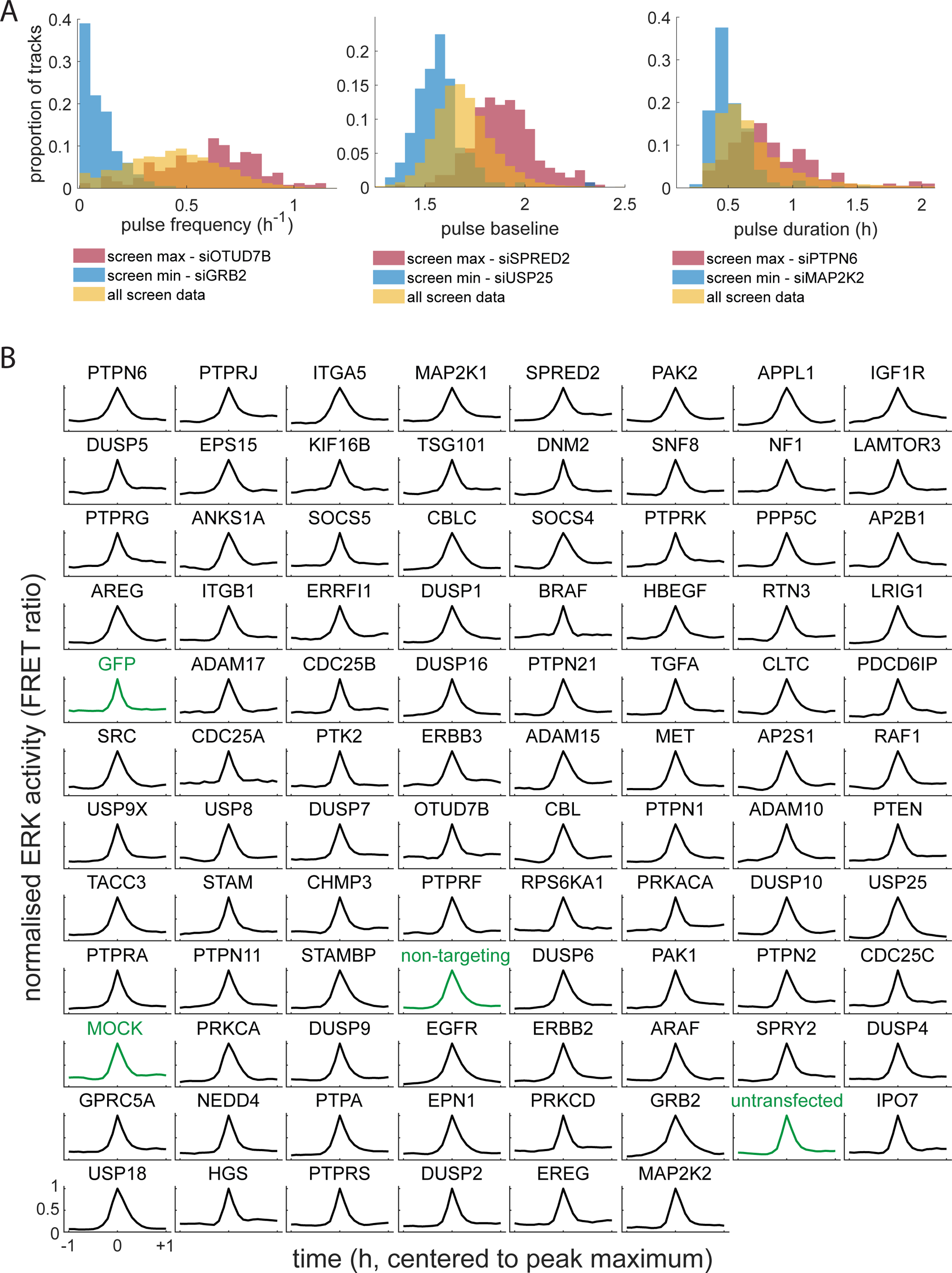
(Related to Fig. 4) (A). Distributions of pulse frequency (left), pulse baseline (middle) and pulse duration (right) in entire screen (yellow) compared to siRNA targets with strongest negative (blue) or positive (red) effects on these metrics. (B). Average normalised pulse shape for all screen targets. Controls are highlighted in green.

**Figure S8.**
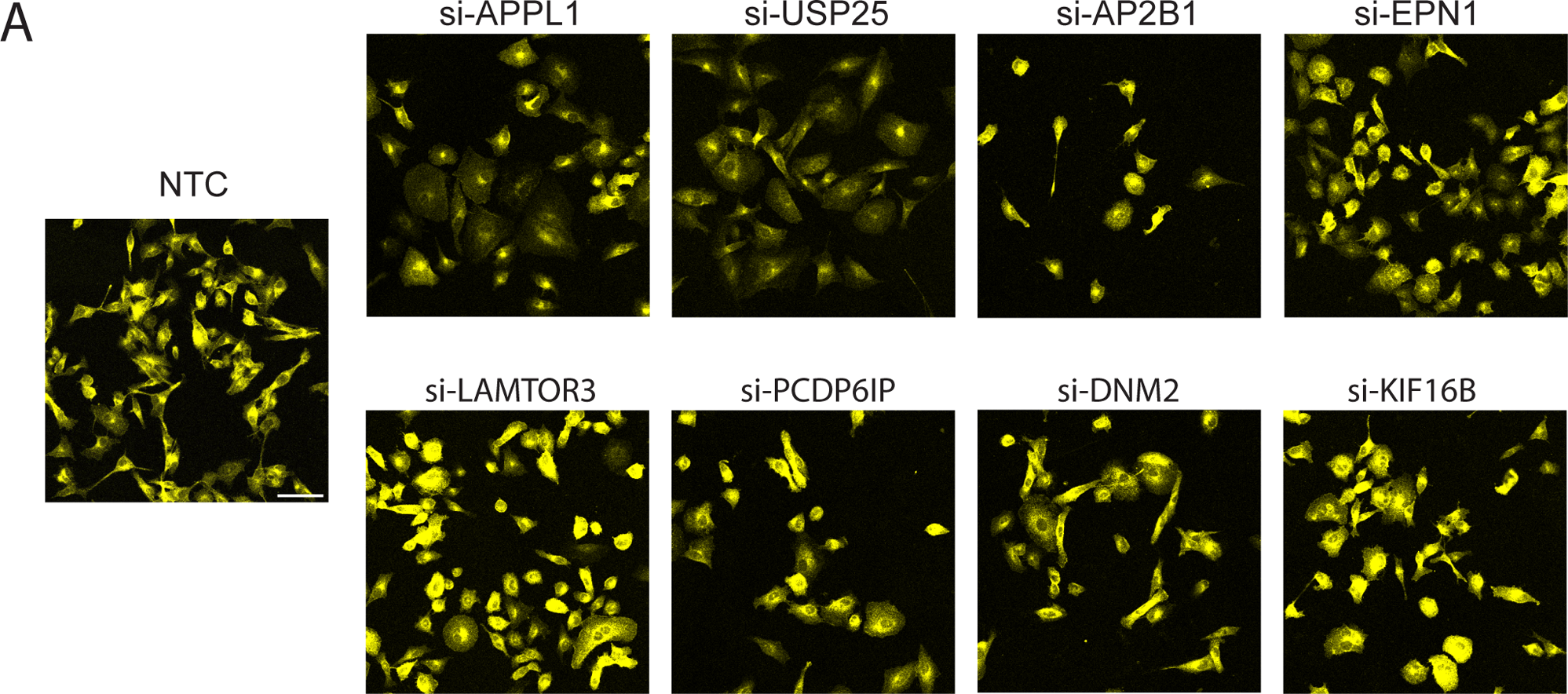
(Related to Fig. 5) (A). Immunofluorescence imaging of EGFR in screen targets involved in membrane trafficking. Scale bar is 100µm.

**Figure S9.**
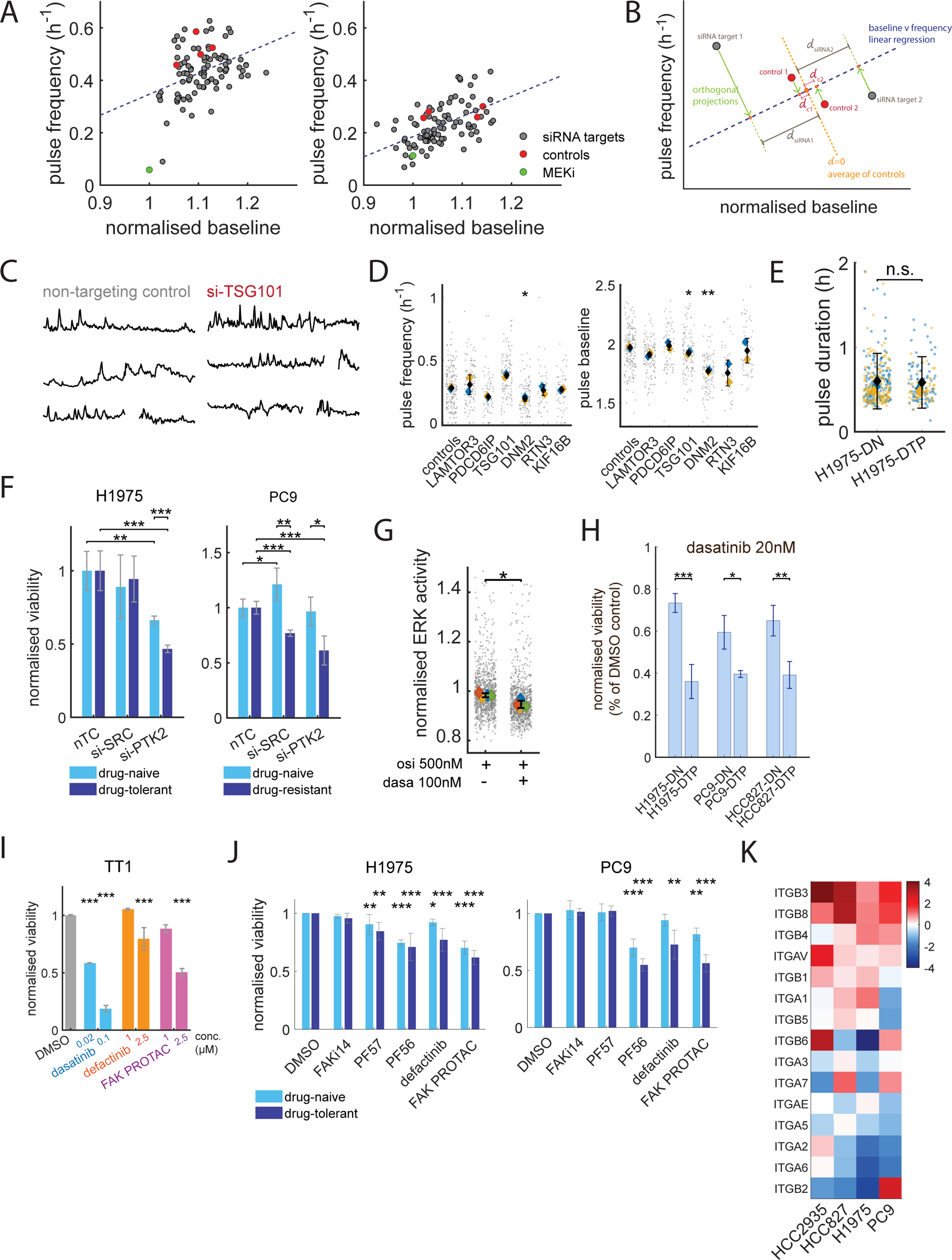
(Related to Fig. 6) (A). ERK pulse baseline and frequency for all the screen targets in drug-naïve (left) and drug-tolerant (right) H1975 cells. The dotted lines represent the line of best fit for linear regression models. (B). Diagram illustrating the method to obtain the ‘distance from control’ metric used to rank the screen targets as shown in Figure 6(A). (C). Example ERK traces for some screen hits. Gaps in traces indicate mitotic events. (D). ERK pulse frequency and baseline with knock down of cluster 3 membrane trafficking genes. (E). Comparison of ERK pulse duration in H1975 drug-naïve and drug-tolerant cells. Data is from N=2 independent experiments. (F). Viability in H1975 drug naïve or osi-DTP’s (left) and PC9 drug naïve or drug-resistant (right) cells upon siRNA-mediated knock-down of SRC or PTK2. Data is from N=3 independent experiments. (G). ERK activity levels upon dasatinib treatment in H1975-DTP cells quantified using the EKAREV-FRET sensor. Data is from N=4 independent experiments. (H). Viability in 3 pairs of drug-naïve and drug-tolerant EGFR-mutant cells lines upon treatment with 20nM dasatinib. Data is from a minimum of N=3 independent experiments. (I). Viability in TT1 cells treated with dasatinib, defactinib or defactinib-derived FAK PROTAC. Data is from N=3 independent experiments. (J). Viability in H1975 and PC9 drug-naïve or drug-tolerant cells treated with a panel of FAK inhibitors. Data is from a minimum of N=3 independent experiments for each condition. (K). Fold-change expression levels of integrin genes in DTPs for 4 cell lines of EGFR-mutant NSCLC, based on the GSE193258 dataset. **Statistical tests:** (D) Unpaired Student’s t-test between the mean of the replicates. (E) Unpaired Student’s t-test between the data points. (F) & (H) Unpaired Student’s t-test between conditions. Error bars show the s.d. of the mean. (G) Unpaired Student’s t-test between the replicate means. (I) & (J). 1-way ANOVA with Tukey’s multiple comparison test.

**Figure S10.**
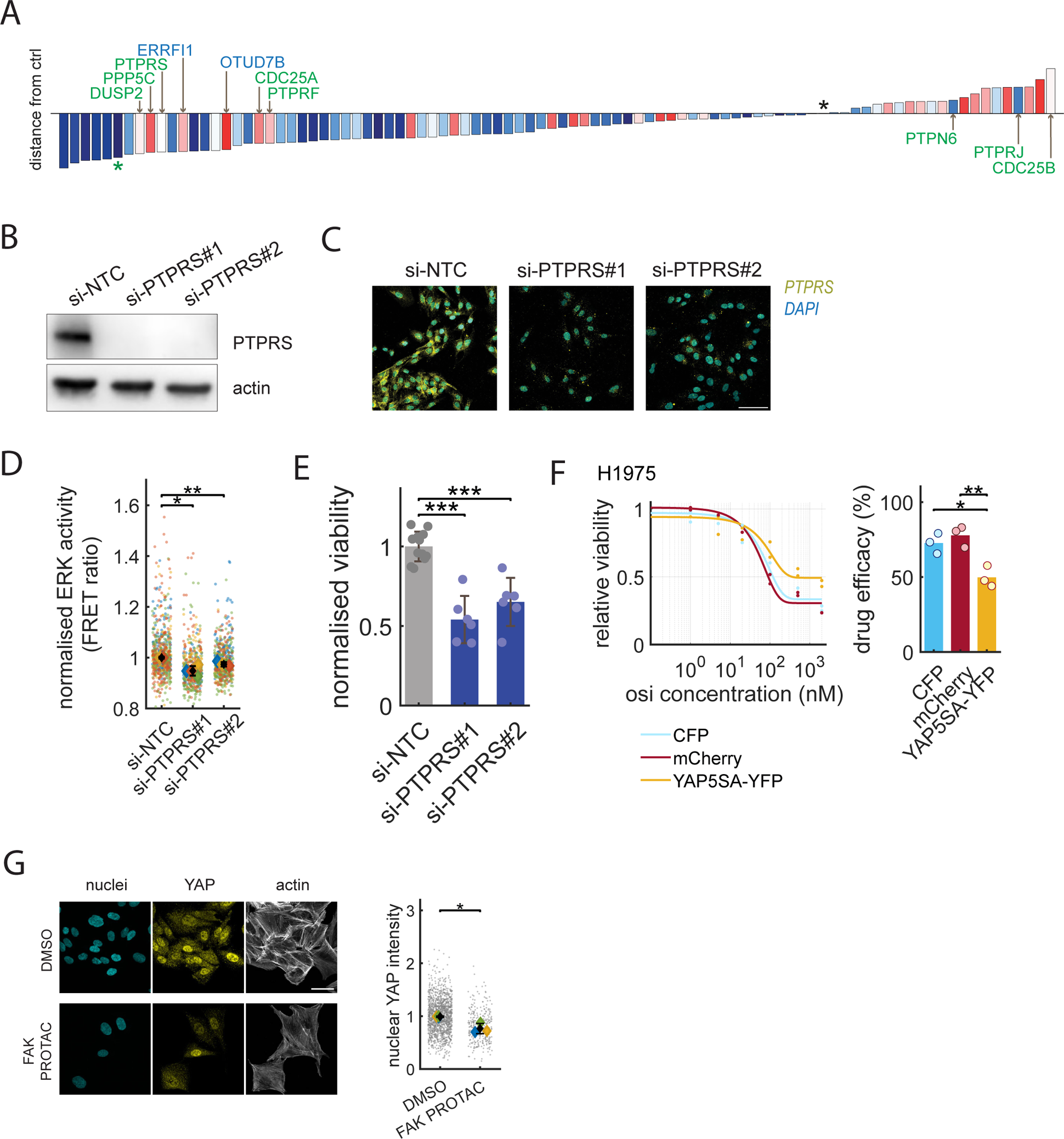
(Related to Fig. 7). (A). Bar charts of the screen targets in the drug-tolerant state ranked in order of distance from the negative controls in terms of their effects on ERK pulse frequency and baseline. Bar colours from the drug-naïve conditions (Fig. 6A) are conserved to highlight targets which are rewired. Data is from N=2 replicates. (B). Effect of siRNA-mediated PTPRS knock-down on PTPRS protein levels detected by immunoblotting. Data is representative of N=2 independent experiments. (C). Effect of siRNA-mediated PTPRS knock-down on PTPRS protein levels detected by immunofluorescence imaging. Data is representative of N=2 independent experiments. (D). Normalised ERK activity levels upon PTPRS knock down. Data is from N=4 independent experiments. (E). Effect of PTPRS knock-down on viability in H1975-DTP cells. Data is from N=6 independent experiments. (F). Osimertinib dose response curves and treatment efficacy quantifications for H1975 cells expressing the YAP5SA-YFP construct or 2 control fluorescent protein constructs (CFP and mCherry). Data is from N=2 independent experiments. (G). Effect of FAK PROTAC treatment on YAP levels. Data is from N=3 independent experiments. Scale bar is 50µm. **Statistical tests:** (D) & (G). Paired Student’s t-test between replicate means. (E) & (F). Unpaired Student’s t-test.

Supplementary Video 1 (related to Fig. 1D).

Illustration of an H1975 cell expressing both ERK (left) and AKT (right) biosensors tracked from one cell division to the next.

Supplementary Video 2 (related to Fig. 1H).

Illustration of ERK and AKT sensor readouts for H1975 cells treated with vehicle control (top row), IGF1R inhibitor (middle), or starvation media (bottom row).

Supplementary Video 3a (related to Fig. 2B-E)

Timelapse imaging of ERK activity (FRET ratio in false colours) in H1975 cells.

Supplementary Video 3b (related to Fig. 2B-E)

Timelapse imaging of ERK activity (FRET ratio in false colours) in TT1 cells.

Supplementary Video 4 (related to Fig. 3A)

Timelapse imaging of ERK activity in PC9 (top row) or H1975 (bottom row) cells, exposed to acute (left column) or chronic (right column) EGFR inhibition using 500nM Osimertinib.

Supplementary Video 5 (related to Fig. S4F)

Illustration of collective ERK activity pulses in PC9-DTPs. Collective event can be seen at 1.7h approx.

Supplementary Video 6 (related to Fig. 4E-F)

Example timelapse images of live-cell siRNA biosensor screen showing ERK activity in control conditions (leftmost panel, MOCK transfection), Cluster 2 screen hits (si-EGFR and si-STAMBP, 2^nd^ and 3^rd^ from left) and Cluster 3 screen hits (si-TSG101 and si-SPRED2, 4^th^ and 5^th^ from left).

## Acknowledgements

The authors acknowledge Paul Smith, former employee of AstraZeneca, and all members of the Sahai lab for their insightful comments on the manuscript. We thank Nathan Curry and Nils Gustafsson from Imperial College London for their work on building the dOPM system at the Francis Crick Institute. **A.L.M.** thanks the Cell Services, Flow Cytometry, Light Microscopy and Biological Research facilities from the Francis Crick Institute for their assistance and training**. A.L.M.** is a recipient of post-doctoral funding by AstraZeneca through the Crick/AZ Alliance. **D.R.C** is supported by the Francis Crick Institute which receives its core funding form Cancer Research UK (FC001169), the UK Medical Research Council (FC002269), and the Wellcome Trust (FC001169), as well as an NC3Rs training fellowship (NC/S001832/1). **K.V.** is supported by a CRUK Accelerator doctoral studentship (C7408/A28450). **E.S.** is supported by the Francis Crick Institute, which receives its core funding from Cancer Research UK (CC2040), the UK Medical Research Council (CC2040), and the Wellcome Trust (CC2040) and the European Research Council (ERC Advanced Grant CAN_ORGANISE, Grant agreement number 101019366). **C.S.** is a Royal Society Napier Research Professor (RSRP\R\210001). His work is supported by the Francis Crick Institute that receives its core funding from Cancer Research UK (CC2041), the UK Medical Research Council (CC2041), and the Wellcome Trust (CC2041) and the European Research Council under the European Union’s Horizon 2020 research and innovation program (ERC Advanced Grant PROTEUS Grant agreement no. 835297). **C.D.** is supported by EPSRC (EP/T003103/1), CRUK Accelerator (C10441/A29368) and Imperial EPSRC Impact Acceleration Account (EP/R511547/1). **P.F.** is supported by a CRUK MDA award (C7408/A28450). **F.F.** is supported by the Francis Crick Institute, which receives its core funding from Cancer Research UK (CC2242), the UK Medical Research Council (CC2242), and the Wellcome Trust (CC2242). **J.D.** is supported by the Francis Crick Institute, which receives its core funding from Cancer Research UK (FC001070), the UK Medical Research Council (FC001070), and the Wellcome Trust (FC001070), the European Research Council (ERC Advanced RASImmune), and from a Wellcome Trust Senior Investigator Award 103799/Z/14/Z.

## Competing interests

**E.S.** reports grants from Novartis, Merck Sharp Dohme, AstraZeneca and personal fees from Phenomic outside the submitted work. **F.F.** receives consulting fees from Deep Origin outside the submitted work. **C.S** reports grants and personal fees from Bristol Myers Squibb, AstraZeneca, Boehringer-Ingelheim, Roche-Ventana, personal fees from Pfizer, grants from Ono Pharmaceutical, Personalis, grants, personal fees, and other support from GRAIL, other support from AstraZeneca and GRAIL, personal fees and other support from Achilles Therapeutics, Bicycle Therapeutics, personal fees from Genentech, Medixci, China Innovation Centre of Roche (CiCoR) formerly Roche Innovation Centre, Metabomed, Relay Therapeutics, Saga Diagnostics, Sarah Canon Research Institute, Amgen, GlaxoSmithKline, Illumina, MSD, Novartis, other support from Apogen Biotechnologies and Epic Bioscience outside the submitted work; in addition, **C.S.** has a patent for PCT/ US2017/028013 licensed to Natera Inc, UCL Business, a patent for PCT/EP2016/059401 licensed to Cancer Research Technology, a patent for PCT/EP2016/071471 issued to Cancer Research Technology, a patent for PCT/GB2018/051912 pending, a patent for PCT/ GB2018/052004 issued to Francis Crick Institute, University College London, Cancer Research Technology Ltd, a patent for PCT/ GB2020/050221 issued to Francis Crick Institute, University College London, a patent for PCT/EP2022/077987 pending to Cancer Research Technology, a patent for PCT/GB2017/053289 licensed, a patent for PCT/EP2022/077987 pending to Francis Crick Institute, a patent for PCT/EP2023/059039 pending to Francis Crick Institute, and a patent for PCT/GB2018/051892 pending to Francis Crick Institute. **C.S** is Co-chief Investigator of NHS Galleri trial funded by GRAIL. He is Chief Investigator for the AstraZeneca MeRmaiD I and II clinical trials and Chair of the Steering Committee. **C.S** is cofounder of Achilles Therapeutics and holds stock options. **M.M.** is an employee and shareholder of AstraZeneca. **C.D.** has a granted patent on oblique plane microscopy (OPM) (PCT/GB2009/001802) licensed to Leica Microsystems and sublicensed to ASI, and has filed a patent application (PCT/GB2020/052279) on dual-view oblique plane microscopy (dOPM). **J.D.** reports grants from Bristol Myers Squibb, Revolution Medicines, Novartis, Vividion, AstraZeneca and personal fees from Jubilant, Theras, Roche, Curve Therapeutics.

---

**Supplementary Table 1:** summary of cell fate and signalling states upon EGFR inhibition in EGFR-mutant NSCLC lines.

| Treatment Status | Drug-naïve | Acute | Chronic (DTP) |
| --- | --- | --- | --- |
| Cell Fates | Fast proliferation<br>No death | Cell cycle arrest<br>Frequent cell death | Slow proliferation<br>Some cell death |
| ERK signalling | Dynamic<br>Frequent pulses | Shut-down<br>No pulses | Dynamic<br>Infrequent pulses |
| PI3K signalling | Sustained<br>Elevated levels | Sustained<br>Cell line-dependent | Sustained<br>Cell line-dependent |
| Cell fate correlated w. ERK signalling ? | Yes | No | Yes |

**Supplementary Table 2.**

Cluster 2 genes
|  |  |
| --- | --- |
| TACC3 | Mitotic spindle localisation |
| EPN1 | Epsin1 - CME effector |
| DUSP16 | Dual-specificity phosphatase |
| ARAF | MAPK pathway member |
| PTPRA | Tyrosine phosphatase |
| ITGA5 | Integrin alpha 5 |
| USP25 | De-ubiquitinating enzyme |
| MET | Receptor Tyrosine Kinase |
| STAMBP | De-ubiquitinating enzyme |
| APPL1 | Early endosome function |
| USP18 | Regulation of EGFR translation |
| PTK2 | FAK - tyrosine kinase |
| AREG | EGFR ligand |
| PTPN11 | SHP2 - RTK signal transduction |
| CBLC | RTK ubiquitination |
| AP2B1 | AP2 subunit - CME effector |
| EGFR | Receptor Tyrosine Kinase |
| GRB2 | RTK signal transduction |

**Supplementary Table 3.**
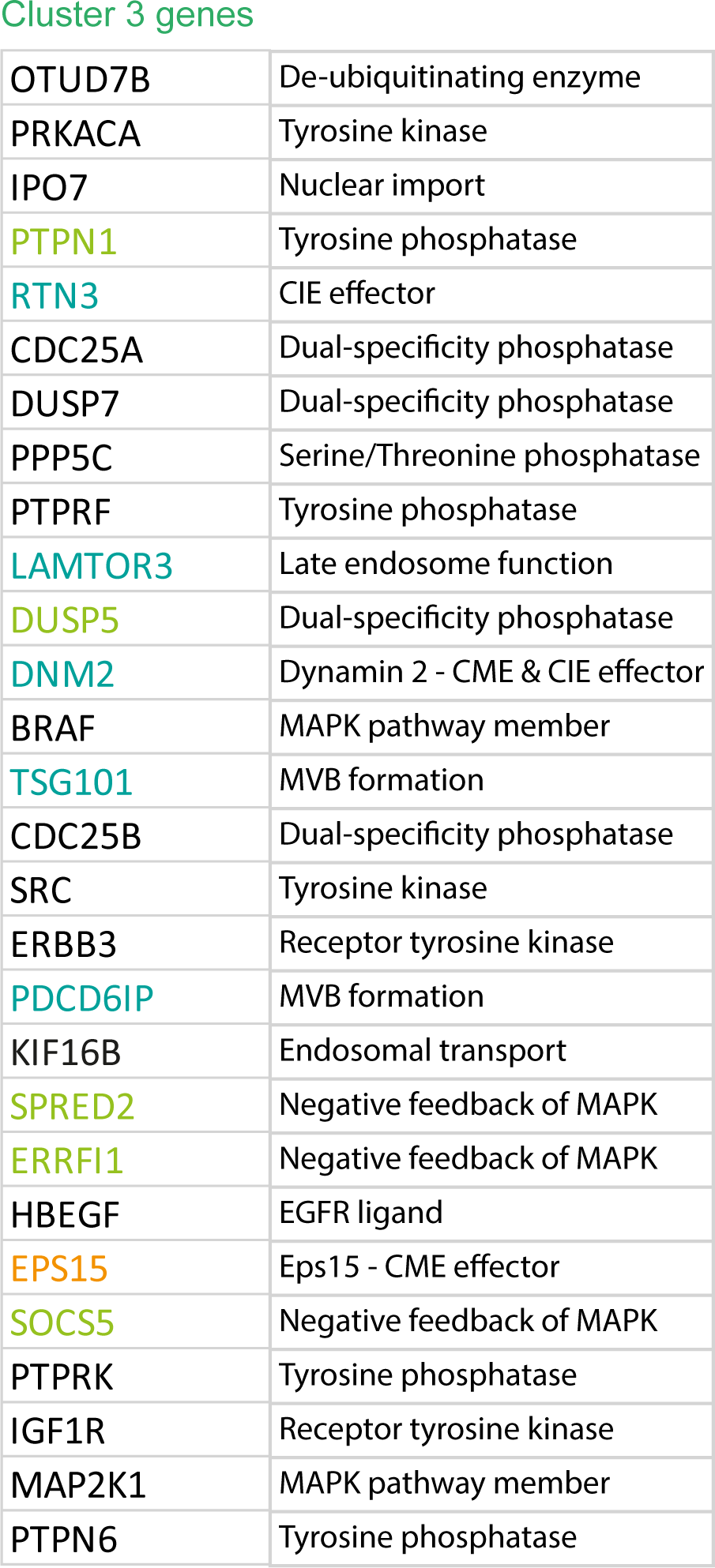

Cluster 3 genes
|  |  |
| --- | --- |
| OTUD7B | De-ubiquitinating enzyme |
| PRKACA | Tyrosine kinase |
| IPO7 | Nuclear import |
| PTPN1 | Tyrosine phosphatase |
| RTN3 | CIE effector |
| CDC25A | Dual-specificity phosphatase |
| DUSP7 | Dual-specificity phosphatase |
| PPP5C | Serine/Threonine phosphatase |
| PTPRF | Tyrosine phosphatase |
| LAMTOR3 | Late endosome function |
| DUSP5 | Dual-specificity phosphatase |
| DNM2 | Dynamin 2 - CME & CIE effector |
| BRAF | MAPK pathway member |
| TSG101 | MVB formation |
| CDC25B | Dual-specificity phosphatase |
| SRC | Tyrosine kinase |
| ERBB3 | Receptor tyrosine kinase |
| PDCD6IP | MVB formation |
| KIF16B | Endosomal transport |
| SPRED2 | Negative feedback of MAPK |
| ERRFI1 | Negative feedback of MAPK |
| HBEGF | EGFR ligand |
| EPS15 | Eps15 - CME effector |
| SOCS5 | Negative feedback of MAPK |
| PTPRK | Tyrosine phosphatase |
| IGF1R | Receptor tyrosine kinase |
| MAP2K1 | MAPK pathway member |
| PTPN6 | Tyrosine phosphatase |

## Notes

### Competing Interest Statement

Acknowledgements
A.L.M. is a recipient of post-doctoral funding by AstraZeneca through the Crick/AZ Alliance. D.R.C is supported by the Francis Crick Institute, which receives its core funding form Cancer Research UK (FC001169), the UK Medical Research Council (FC002269), and the Wellcome Trust (FC001169), as well as an NC3Rs training fellowship (NC/S001832/1). K.V. is supported by a CRUK Accelerator doctoral studentship (C7408/A28450). E.S. is supported by the Francis Crick Institute, which receives its core funding from Cancer Research UK (CC2040), the UK Medical Research Council (CC2040), and the Wellcome Trust (CC2040) and the European Research Council (ERC Advanced Grant CAN_ORGANISE, Grant agreement number 101019366). C.S. is a Royal Society Napier Research Professor (RSRP\R\210001). His work is supported by the Francis Crick Institute that receives its core funding from Cancer Research UK (CC2041), the UK Medical Research Council (CC2041), and the Wellcome Trust (CC2041) and the European Research Council under the European Union Horizon 2020 research and innovation program (ERC Advanced Grant PROTEUS Grant agreement no. 835297). C.D. is supported by EPSRC (EP/T003103/1), CRUK Accelerator (C10441/A29368) and Imperial EPSRC Impact Acceleration Account (EP/R511547/1). P.F. is supported by a CRUK MDA award (C7408/A28450). F.F. is supported by the Francis Crick Institute, which receives its core funding from Cancer Research UK (CC2242), the UK Medical Research Council (CC2242), and the Wellcome Trust (CC2242). J.D. is supported by the Francis Crick Institute, which receives its core funding from Cancer Research UK (FC001070), the UK Medical Research Council (FC001070), and the Wellcome Trust (FC001070), the European Research Council (ERC Advanced RASImmune), and from a Wellcome Trust Senior Investigator Award 103799/Z/14/Z.
Competing interests
E.S. reports grants from Novartis, Merck Sharp Dohme, AstraZeneca and personal fees from Phenomic outside the submitted work. F.F. receives consulting fees from Deep Origin outside the submitted work. C.S reports grants and personal fees from Bristol Myers Squibb, AstraZeneca, BoehringerIngelheim, Roche-Ventana, personal fees from Pfizer, grants from Ono Pharmaceutical, Personalis, grants, personal fees, and other support from GRAIL, other support from AstraZeneca and GRAIL, personal fees and other support from Achilles Therapeutics, Bicycle Therapeutics, personal fees from Genentech, Medixci, China Innovation Centre of Roche (CiCoR) formerly Roche Innovation Centre, Metabomed, Relay Therapeutics, Saga Diagnostics, Sarah Canon Research Institute, Amgen, GlaxoSmithKline, Illumina, MSD, Novartis, other support from Apogen Biotechnologies and Epic Bioscience outside the submitted work; in addition, C.S. has a patent for PCT/ US2017/028013 licensed to Natera Inc, UCL Business, a patent for PCT/EP2016/059401 licensed to Cancer Research Technology, a patent for PCT/EP2016/071471 issued to Cancer Research Technology, a patent for PCT/GB2018/051912 pending, a patent for PCT/ GB2018/052004 issued to Francis Crick Institute, University College London, Cancer Research Technology Ltd, a patent for PCT/ GB2020/050221 issued to Francis Crick Institute, University College London, a patent for PCT/EP2022/077987 pending to Cancer Research Technology, a patent for PCT/GB2017/053289 licensed, a patent for PCT/EP2022/077987 pending to Francis Crick Institute, a patent for PCT/EP2023/059039 pending to Francis Crick Institute, and a patent for PCT/GB2018/051892 pending to Francis Crick Institute. C.S is Co-chief Investigator of NHS Galleri trial funded by GRAIL. He is Chief Investigator for the AstraZeneca MeRmaiD I and II clinical trials and Chair of the Steering Committee. C.S is cofounder of Achilles Therapeutics and holds stock options. M.M. is an employee and shareholder of AstraZeneca. C.D. has a granted patent on oblique plane microscopy (OPM) (PCT/GB2009/001802) licensed to Leica Microsystems and sublicensed to ASI, and has filed a patent application (PCT/GB2020/052279) on dual-view oblique plane microscopy (dOPM). J.D. reports grants from Bristol Myers Squibb, Revolution Medicines, Novartis, Vividion, AstraZeneca and personal fees from Jubilant, Theras, Roche, Curve Therapeutics.

## References

1. Ding, L. et al. Somatic mutations affect key pathways in lung adenocarcinoma. Nature 455, 1069–1075 (2008).

2. Seo, J.-S. et al. The transcriptional landscape and mutational profile of lung adenocarcinoma. Genome Res. 22, 2109–2119 (2012).

3. Wee, P. & Wang, Z. Epidermal Growth Factor Receptor Cell Proliferation Signaling Pathways. Cancers 9, 52 (2017).

4. Liu, J., Kang, R. & Tang, D. The KRAS-G12C inhibitor: activity and resistance. Cancer Gene Ther. 29, 875–878 (2022).

5. Cataldo Vince D., Gibbons Don L., Pérez-Soler Román, & Quintás-Cardama Alfonso. Treatment of Non–Small-Cell Lung Cancer with Erlotinib or Gefitinib. N. Engl. J. Med. 364, 947–955 (2011).

6. Ramalingam Suresh S., et al. Overall Survival with Osimertinib in Untreated, EGFR-Mutated Advanced NSCLC. N. Engl. J. Med. 382, 41–50 (2020).

7. Canon, J. et al. The clinical KRAS(G12C) inhibitor AMG 510 drives anti-tumour immunity. Nature 575, 217–223 (2019).

8. Sharma, S. V. et al. A Chromatin-Mediated Reversible Drug-Tolerant State in Cancer Cell Subpopulations. Cell 141, 69–80 (2010).

9. Russo, M. et al. Adaptive mutability of colorectal cancers in response to targeted therapies. Science 366, 1473–1480 (2019).

10. Oren, Y. et al. Cycling cancer persister cells arise from lineages with distinct programs. Nature 596, 576–582 (2021).

11. Lake, D., Corrêa, S. A. L. & Müller, J. Negative feedback regulation of the ERK1/2 MAPK pathway. Cell. Mol. Life Sci. 73, 4397–4413 (2016).

12. Tomas, A., Futter, C. E. & Eden, E. R. EGF receptor trafficking: consequences for signaling and cancer. Trends Cell Biol. 24, 26–34 (2014).

13. Polo, S., Di Fiore, P. P. & Sigismund, S. Keeping EGFR signaling in check: Ubiquitin is the guardian. Cell Cycle 13, 681–682 (2014).

14. Sigismund, S. et al. Threshold-controlled ubiquitination of the EGFR directs receptor fate. EMBO J. 32, 2140–2157 (2013).

15. Hong, S.-Y., Kao, Y.-R., Lee, T.-C. & Wu, C.-W. Upregulation of E3 Ubiquitin Ligase CBLC Enhances EGFR Dysregulation and Signaling in Lung Adenocarcinoma. Cancer Res. 78, 4984–4996 (2018).

16. Purvis, J. E. et al. p53 dynamics control cell fate. Science 336, 1440–1444 (2012).

17. Aoki, K. et al. Propagating Wave of ERK Activation Orients Collective Cell Migration. Dev. Cell 43, 305–317.e5 (2017).

18. Hiratsuka, T., Bordeu, I., Pruessner, G. & Watt, F. M. Regulation of ERK basal and pulsatile activity control proliferation and exit from the stem cell compartment in mammalian epidermis. Proc. Natl. Acad. Sci. 117, 17796–17807 (2020).

19. Gagliardi, P. A. et al. Collective ERK/Akt activity waves orchestrate epithelial homeostasis by driving apoptosis-induced survival. Dev. Cell 56, 1712–1726.e6 (2021).

20. Albeck, J. G., Mills, G. B. & Brugge, J. S. Frequency-Modulated Pulses of ERK Activity Transmit Quantitative Proliferation Signals. Mol. Cell 49, 249–261 (2013).

21. Goglia, A. G. et al. A Live-Cell Screen for Altered Erk Dynamics Reveals Principles of Proliferative Control. Cell Syst. 10, 240–253.e6 (2020).

22. Komatsu, N. et al. Development of an optimized backbone of FRET biosensors for kinases and GTPases. Mol. Biol. Cell 22, 4647–4656 (2011).

23. Kudo, T. et al. Live-cell measurements of kinase activity in single cells using translocation reporters. Nat. Protoc. 13, 155–169 (2018).

24. Kemp, S. J. et al. Immortalization of Human Alveolar Epithelial Cells to Investigate Nanoparticle Uptake. Am. J. Respir. Cell Mol. Biol. 39, 591–597 (2008).

25. Ponsioen, B. et al. Quantifying single-cell ERK dynamics in colorectal cancer organoids reveals EGFR as an amplifier of oncogenic MAPK pathway signalling. Nat. Cell Biol. 23, 377–390 (2021).

26. Karachaliou, N. et al. Common Co-activation of AXL and CDCP1 in EGFR-mutation-positive Non-Small Cell Lung Cancer Associated With Poor Prognosis. EBioMedicine 29, 112–127 (2018).

27. Gerosa, L. et al. Receptor-Driven ERK Pulses Reconfigure MAPK Signaling and Enable Persistence of Drug-Adapted BRAF-Mutant Melanoma Cells. Cell Syst. 11, 478–494.e9 (2020).

28. Gagliardi, P. A. et al. Automatic detection of spatio-temporal signaling patterns in cell collectives. J. Cell Biol. 222, e202207048 (2023).

29. Sparks, H. et al. Dual-view oblique plane microscopy (dOPM). Biomed. Opt. Express 11, 7204–7220 (2020).

30. Klijn, C. et al. A comprehensive transcriptional portrait of human cancer cell lines. Nat. Biotechnol. 33, 306–312 (2015).

31. Barretina, J. et al. The Cancer Cell Line Encyclopedia enables predictive modelling of anticancer drug sensitivity. Nature 483, 603–607 (2012).

32. Ghandi, M. et al. Next-generation characterization of the Cancer Cell Line Encyclopedia. Nature 569, 503–508 (2019).

33. Cairns, J. et al. Differential roles of ERRFI1 in EGFR and AKT pathway regulation affect cancer proliferation. EMBO Rep. 19, e44767 (2018).

34. Haj, F. G., Markova, B., Klaman, L. D., Bohmer, F. D. & Neel, B. G. Regulation of Receptor Tyrosine Kinase Signaling by Protein Tyrosine Phosphatase-1B*. J. Biol. Chem. 278, 739–744 (2003).

35. Wakioka, T. et al. Spred is a Sprouty-related suppressor of Ras signalling. Nature 412, 647–651 (2001).

36. Diggins, N. L. & Webb, D. J. APPL1 is a multifunctional endosomal signaling adaptor protein. Biochem. Soc. Trans. 45, 771–779 (2017).

37. Sorkin, A. & von Zastrow, M. Endocytosis and signalling: intertwining molecular networks. Nat. Rev. Mol. Cell Biol. 10, 609–622 (2009).

38. Monypenny, J. et al. ALIX Regulates Tumor-Mediated Immunosuppression by Controlling EGFR Activity and PD-L1 Presentation. Cell Rep. 24, 630–641 (2018).

39. Caldieri, G. et al. Reticulon 3-dependent ER-PM contact sites control EGFR nonclathrin endocytosis. Science 356, 617–624 (2017).

40. Nagy, Z., Mori, J., Ivanova, V.-S., Mazharian, A. & Senis, Y. A. Interplay between the tyrosine kinases Chk and Csk and phosphatase PTPRJ is critical for regulating platelets in mice. Blood 135, 1574–1587 (2020).

41. Ichihara, E. et al. SFK/FAK Signaling Attenuates Osimertinib Efficacy in Both Drug-Sensitive and Drug-Resistant Models of EGFR-Mutant Lung Cancer. Cancer Res. 77, 2990–3000 (2017).

42. Creelan, B. C. et al. Phase 1 trial of dasatinib combined with afatinib for epidermal growth factor receptor-(EGFR-) mutated lung cancer with acquired tyrosine kinase inhibitor (TKI) resistance. Br. J. Cancer 120, 791–796 (2019).

43. Johnson, M. L. et al. Phase II Trial of Dasatinib for Patients with Acquired Resistance to Treatment with the Epidermal Growth Factor Receptor Tyrosine Kinase Inhibitors Erlotinib or Gefitinib. J. Thorac. Oncol. 6, 1128–1131 (2011).

44. Kim, C. et al. A Phase I Trial of Dasatinib and Osimertinib in TKI Naïve Patients With Advanced EGFR-Mutant Non-Small-Cell Lung Cancer. Front. Oncol. 11, 728155 (2021).

45. Cromm, P. M., Samarasinghe, K. T. G., Hines, J. & Crews, C. M. Addressing Kinase-Independent Functions of Fak via PROTAC-Mediated Degradation. J. Am. Chem. Soc. 140, 17019–17026 (2018).

46. Gao, H. et al. FAK-targeting PROTAC as a chemical tool for the investigation of non-enzymatic FAK function in mice. Protein Cell 11, 534–539 (2020).

47. Mitra, S. K. & Schlaepfer, D. D. Integrin-regulated FAK–Src signaling in normal and cancer cells. Curr. Opin. Cell Biol. 18, 516–523 (2006).

48. Kawauchi, T. Cell Adhesion and Its Endocytic Regulation in Cell Migration during Neural Development and Cancer Metastasis. Int. J. Mol. Sci. 13, 4564–4590 (2012).

49. Moreno-Layseca, P., Icha, J., Hamidi, H. & Ivaska, J. Integrin trafficking in cells and tissues. Nat. Cell Biol. 21, 122–132 (2019).

50. Alanko, J. et al. Integrin endosomal signalling suppresses anoikis. Nat. Cell Biol. 17, 1412–1421 (2015).

51. Gogleva, A. et al. Knowledge graph-based recommendation framework identifies drivers of resistance in EGFR mutant non-small cell lung cancer. Nat. Commun. 13, 1667 (2022).

52. Looyenga, B. D. & MacKeigan, J. P. Characterization of Differential Protein Tethering at the Plasma Membrane in Response to Epidermal Growth Factor Signaling. J. Proteome Res. 11, 3101–3111 (2012).

53. Mu, F.-T. et al. EEA1, an Early Endosome-Associated Protein. J. Biol. Chem. 270, 13503–13511 (1995).

54. Pocaterra, A., Romani, P. & Dupont, S. YAP/TAZ functions and their regulation at a glance. J. Cell Sci. 133, jcs230425 (2020).

55. Calvo, F. et al. Mechanotransduction and YAP-dependent matrix remodelling is required for the generation and maintenance of cancer-associated fibroblasts. Nat. Cell Biol. 15, 637–646 (2013).

56. Kurppa, K. J. et al. Treatment-Induced Tumor Dormancy through YAP-Mediated Transcriptional Reprogramming of the Apoptotic Pathway. Cancer Cell 37, 104–122.e12 (2020).

57. Ege, N. et al. Quantitative Analysis Reveals that Actin and Src-Family Kinases Regulate Nuclear YAP1 and Its Export. Cell Syst. 6, 692–708.e13 (2018).

58. Pinilla-Macua, I., Watkins, S. C. & Sorkin, A. Endocytosis separates EGF receptors from endogenous fluorescently labeled HRas and diminishes receptor signaling to MAP kinases in endosomes. Proc. Natl. Acad. Sci. 113, 2122–2127 (2016).

59. Surve, S., Watkins, S. C. & Sorkin, A. EGFR-RAS-MAPK signaling is confined to the plasma membrane and associated endorecycling protrusions. J. Cell Biol. 220, e202107103 (2021).

60. Villaseñor, R., Nonaka, H., Del Conte-Zerial, P., Kalaidzidis, Y. & Zerial, M. Regulation of EGFR signal transduction by analogue-to-digital conversion in endosomes. eLife 4, e06156 (2015).

61. Fortian, A. & Sorkin, A. Live-cell fluorescence imaging reveals high stoichiometry of Grb2 binding to the EGF receptor sustained during endocytosis. J. Cell Sci. 127, 432–444 (2014).

62. Villaseñor, R., Kalaidzidis, Y. & Zerial, M. Signal processing by the endosomal system. Curr. Opin. Cell Biol. 39, 53–60 (2016).

63. Davis, T. B. et al. PTPRS Regulates Colorectal Cancer RAS Pathway Activity by Inactivating Erk and Preventing Its Nuclear Translocation. Sci. Rep. 8, 9296 (2018).

64. Wang, Z.-C. et al. Protein tyrosine phosphatase receptor S acts as a metastatic suppressor in hepatocellular carcinoma by control of epithermal growth factor receptor–induced epithelial-mesenchymal transition. Hepatology 62, 1201–1214 (2015).

65. Morris, L. G. T. et al. Genomic dissection of the epidermal growth factor receptor (EGFR)/PI3K pathway reveals frequent deletion of the EGFR phosphatase PTPRS in head and neck cancers. Proc. Natl. Acad. Sci. U. S. A. 108, 19024–19029 (2011).

66. Davis, T. B. et al. PTPRS drives adaptive resistance to MEK/ERK inhibitors through SRC. Oncotarget 10, 6768–6780 (2019).

67. Cornejo, F., Cortés, B. I., Findlay, G. M. & Cancino, G. I. LAR Receptor Tyrosine Phosphatase Family in Healthy and Diseased Brain. Front. Cell Dev. Biol. 9, (2021).

68. Caswell, D. R. et al. The role of APOBEC3B in lung tumor evolution and targeted cancer therapy resistance. Nat. Genet. 56, 60–73 (2024).

69. Hirata, E. et al. Intravital Imaging Reveals How BRAF Inhibition Generates Drug-Tolerant Microenvironments with High Integrin β1/FAK Signaling. Cancer Cell 27, 574–588 (2015).

70. Gross, S. M. & Rotwein, P. Akt signaling dynamics in individual cells. J. Cell Sci. 128, 2509–2519 (2015).

71. Ershov, D. et al. TrackMate 7: integrating state-of-the-art segmentation algorithms into tracking pipelines. Nat. Methods 19, 829–832 (2022).

72. Yang, H. W., Chung, M., Kudo, T. & Meyer, T. Competing memories of mitogen and p53 signalling control cell-cycle entry. Nature 549, 404–408 (2017).

73. Margineanu, A. et al. Screening for protein-protein interactions using Förster resonance energy transfer (FRET) and fluorescence lifetime imaging microscopy (FLIM). Sci. Rep. 6, 28186 (2016).

74. Alexandrov, Y. et al. Method and software for model-based artifact correction in time-lapse fluorescence microscopy images. in (The Francis Crick Institute, 2022). doi:10.25418/crick.19626810.v1.

75. Akram, K. M. et al. Live imaging of alveologenesis in precision-cut lung slices reveals dynamic epithelial cell behaviour. Nat. Commun. 10, 1178 (2019).

76. Dunsby, C. Optically sectioned imaging by oblique plane microscopy. Opt. Express 16, 20306–20316 (2008).

77. Smith, M. B. et al. Active mesh and neural network pipeline for cell aggregate segmentation. Biophys. J. 122, 1586–1599 (2023).

78. Preibisch, S. et al. Efficient Bayesian-based multiview deconvolution. Nat. Methods 11, 645–648 (2014).

79. Santos, A. F., et al. Angiogenesis: An Improved In Vitro Biological System and Automated Image-Based Workflow to Aid Identification and Characterization of Angiogenesis and Angiogenic Modulators. ASSAY Drug Dev. Technol. 6, 693–710 (2008).

80. Guglielmi, L. et al. Smad4 controls signaling robustness and morphogenesis by differentially contributing to the Nodal and BMP pathways. Nat. Commun. 12, 6374 (2021).

81. Nguyen, T. T. et al. Robust dose-response curve estimation applied to high content screening data analysis. Source Code Biol. Med. 9, 27 (2014).

82. Ge, S. X., Jung, D. & Yao, R. ShinyGO: a graphical gene-set enrichment tool for animals and plants. Bioinformatics 36, 2628–2629 (2020).

83. Szklarczyk, D. et al. The STRING database in 2023: protein-protein association networks and functional enrichment analyses for any sequenced genome of interest. Nucleic Acids Res. 51, D638–D646 (2023).

84. Shannon, P. et al. Cytoscape: a software environment for integrated models of biomolecular interaction networks. Genome Res. 13, 2498–2504 (2003).

